# Repeated experience drives multiscale engram reorganization to shape memory strength and precision

**DOI:** 10.64898/2026.02.01.703091

**Authors:** Mujun Kim, Boin Suh, Yire Jeong, Sunhoi So, Jung Wook Choi, Jaemin Hwang, Juhee Park, Jin-Hee Han

## Abstract

How memories are shaped by repeated experiences remains unclear. Reorganization of memory engram could underlie memory transformation. Here, by combining c-Fos-based engram cell tagging with optogenetics, spine imaging, and reactivation analyses in the dentate gyrus (DG) and medial prefrontal cortex (mPFC), we discovered a distinct system-wide engram reorganization following repeated learning; that is, a reshuffling of engram cells in the DG, as well as the rapid maturation of mPFC engram cells, forming a dual engram distributed within the hippocampal-mPFC network. These mPFC engram cells were crucial for relearning-dependent enhancement of memory precision. In addition, we found that reactivation of DG engram cells during retraining is critical for engram reorganization-dependent memory strengthening. These findings establish a causal link between engram reorganization and memory transformation by relearning.

## INTRODUCTION

Our daily lives mostly consist of repeated experiences. Once formed, memories are continually reshaped by subsequent experiences, leading to the transformation of memory, e.g., changes in memory strength and precision^1,2,3,4^. However, the underlying mechanism remains largely unknown.

Memories are thought to be encoded and stored in a specific group of cells in the brain, called engram cells^5^. Studies of memory engram cells have provided insight into the mechanisms of memory encoding and storage ^5,6,7,8^. Techniques that allow selective labeling and manipulation of cells active during an event have identified neurons expressing immediate-early genes (IEGs), such as c-Fos and Arc (activity-regulated cytoskeleton-associated protein), as engram cells^9,10,11,12^. Recall of a specific memory requires the neurons that were active during its encoding. Recent studies show that memory engram cells and their connectivity are not static but dynamically reorganized by experiences and time^13,14,15,16,17,18,19,20^. For instance, repeated context exposure reveals day-to-day turnover in the population of CA1 pyramidal neurons, as shown by Ca2+ imaging or in vivo electrophysiological recordings^21,22^.We previously found that repeated associative learning induces a turnover of cell populations in the amygdala that supports fear memory^16^. After repeated exposure, initial memory engram cells become dispensable for recall and are replaced by cells activated during retraining. While this study shows that engram cells are reorganized by repeated learning in specific brain areas, such as the amygdala, it remains unclear what happens in broader brain networks. In addition, there have been conflicting views^23,24,25^.

The hippocampal-mPFC networks are essential for encoding, consolidation, and storing episodic memories. In both regions, engram cells have been identified^11,26,27,28,29,30^. The standard model of memory consolidation posits that memory is initially encoded rapidly in the hippocampus and then gradually transferred to the neocortical networks for long-term storage through a process called the systems consolidation. Using engram cell tagging methods, previous studies have demonstrated a time-dependent dynamic reorganization of engram cells and their connections as well as the stable components of the engram within the hippocampal-mPFC networks^18,28,29,31^. Unlike the standard model, it has also been demonstrated that mPFC engram cells form rapidly during memory encoding but mature progressively over time and are reactivated during remote memory retrieval^28,32^.

These findings suggest that the dynamic reorganization of engram cells could underlie memory transformation. Supporting this, recent studies have shown that memory transformation correlates with the dynamic reorganization of engram cells^17,18,19,20^. However, the relationship between engram reorganization and memory transformation is unclear. Current evidence is mainly correlational, and most studies have focused on engram reorganization without directly demonstrating the functional contribution of reorganized engram cells or circuits to changes in memory, such as memory strengthening and precision. Therefore, how an engram is reorganized by repeated experiences at the systems level, and the direct causal link between this reorganization and memory transformation remains obscure.

To address these questions, here we tracked engram cells across repeated learning by combining a c-Fos-based engram cell tagging with optogenetics, spine imaging, and reactivation analyses in the DG and mPFC of transgenic mice^29,33^.

## RESULTS

### c-Fos ensembles in the dentate gyrus are required for contextual fear memory retrieval

To label and manipulate DG neurons activated during memory encoding, we employed FosTRAP mice, which express a tamoxifen-dependent Cre recombinase fused to the estrogen receptor triple mutant 2 (CreERT2) under the control of the c-Fos promoter^33^. In the absence of tamoxifen, CreERT2 remains inactive in the cytoplasm. Upon administration of 4-hydroxytamoxifen (4-OHT), CreERT2 translocates to the nucleus and drives recombination of Cre-dependent AAV vectors in recently active neurons (Figure S1A). To validate the inducible and learning-dependent cell labeling, we injected AAV-EF1α-DIO-EYFP into the bilateral DG of FosTRAP mice and examined EYFP expression under different treatments (Figure S1B). Mice that received 4-OHT immediately after contextual fear conditioning showed significantly higher EYFP+ cells compared to those that either remained in the home cage prior to 4-OHT or received vehicle after training (Figures S1C and S1D), validating that the labeling depends on both 4-OHT administration and learning. We next examined the identity of the labeled neurons. EYFP+ cells were predominantly localized in the granule cell layer rather than the hilus (Figures S2A and S2B). Immunohistochemistry revealed that these EYFP+ cells did not colocalize with GAD67+ cells, indicating that the 4-OHT and learning-dependent labeled neurons were primarily excitatory granule cells (Figure S2C).

Next, FosTRAP mice received bilateral DG injection of AAV-EF1α-DIO-eNpHR3.0-EYFP to label neurons activated during single-shock contextual fear conditioning (Training,1x) with NpHR (Figures 1A and 1B). Two days later, during the memory retrieval test, these labeled neurons were optogenetically silenced using 561 nm light. Mice in the light illuminated group (ON group) displayed significantly reduced freezing behavior to the conditioned context during light exposure compared to the control mice (Figures 1C and 1D). Notably, even when the number of pairings was increased to 2x during training, the ON group continued to show a reduction in freezing behavior compared to controls (Figures 1E and 1F). To verify that NpHR-mediated inhibition specifically disrupted contextual fear memory, we applied the same optogenetic silencing in an auditory fear conditioning paradigm. Inhibiting DG engram cells during tone presentation had no effect on tone-induced freezing but significantly reduced freezing in the conditioned context, demonstrating that the manipulation selectively disrupted memory retrieval rather than freezing behavior per se (Figures 1G-1I). Together, these results demonstrate that DG engram cells are essential for the retrieval of contextual fear memories.

**Figure 1.**
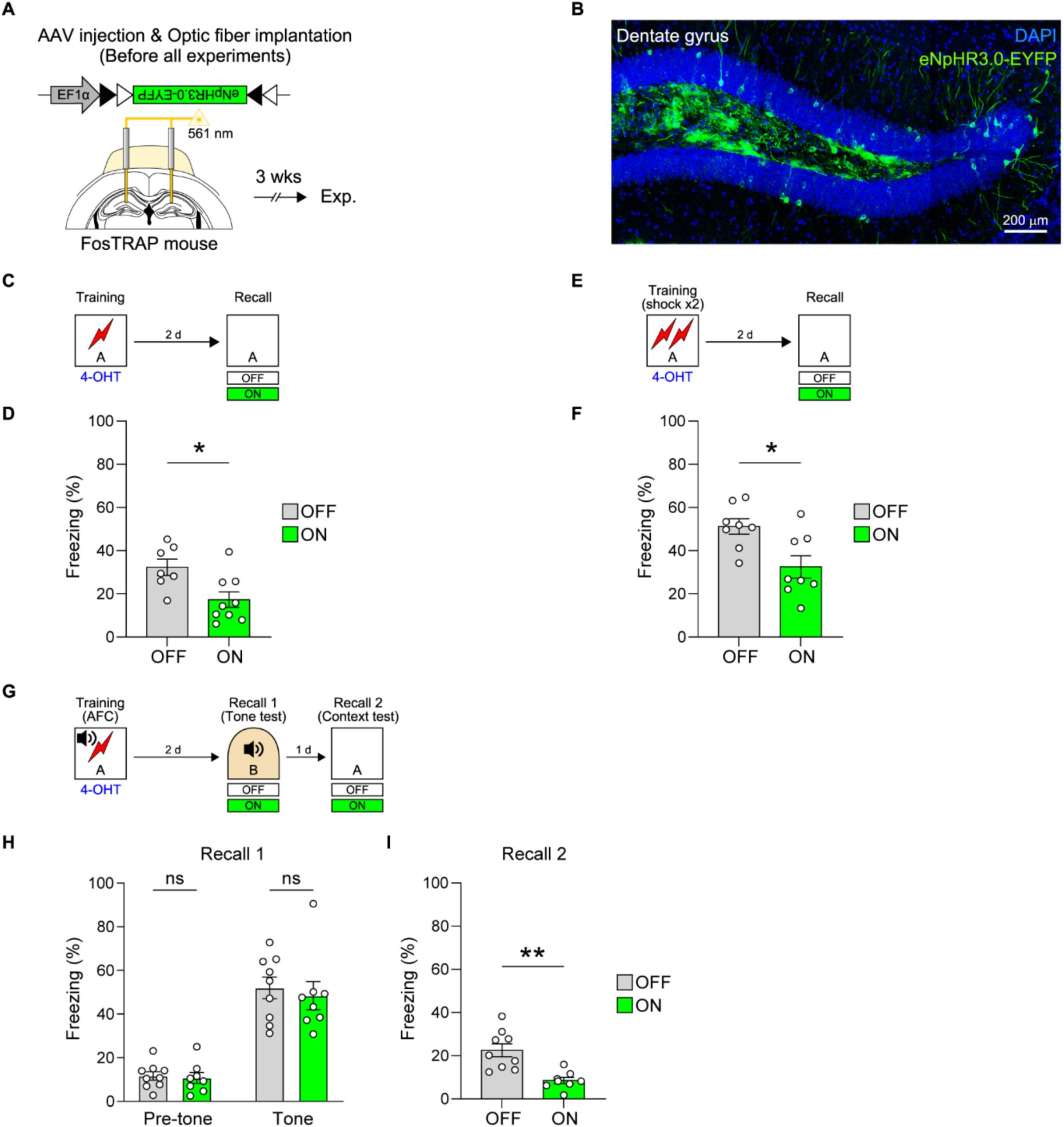
c-Fos ensembles in the dentate gyrus are required for memory retrieval. (A) Schematic of surgery. AAV1-EF1α-DIO-eNpHR3.0-EYFP was bilaterally injected into the DG of FosTRAP mice, followed by implantation of optic fibers above the injection sites. Behavioral experiments were conducted three weeks later. (B) Representative confocal image showing EYFP expression in the DG following training and 4-OHT administration. Scale bar, 200 μm. (C) Behavioral procedure. Mice underwent training, followed by 4-OHT injection immediately after training to label c-Fos ensembles with NpHR-EYFP. During the recall test, the ON group received 561 nm laser illumination, while the OFF group was tested without light (OFF group, n=7 mice; ON group, n=9 mice). (D) Quantification of freezing behavior during the recall test. Optogenetic inhibition of c-Fos ensembles reduced freezing in the ON group compared to the OFF group (p = 0.0131, unpaired t-test). (E) Behavioral procedure. Mice underwent training with two context-shock (2x) pairings, followed by 4-OHT injection to label c-Fos ensembles. During the recall test, the ON group received light illumination, while the OFF group was tested without light (OFF group, n=8 mice; ON group, n=8 mice). (F) Quantification of freezing behavior during the recall test. Optogenetic inhibition of c-Fos ensembles reduced freezing in the ON group compared to the OFF group (p = 0.0105, unpaired t-test). (G) Behavioral procedure. Mice underwent auditory fear conditioning, followed by 4-OHT injection to label c-Fos ensembles. During both the tone and context recall tests, the ON group received light illumination, while the OFF group was tested without light (OFF group, n=9 mice; ON group, n=8 mice). (H) Quantification of freezing behavior during the tone recall test. Freezing was measured during the 3-minute pre-tone period and the 90-second tone period. Light illumination was applied during tone period. Both groups displayed increased freezing during the tone period compared to the pre-tone period (OFF: Pre-tone vs Tone, p<0.0001; ON: Pre-tone vs Tone, p<0.0001). No difference in freezing was observed between OFF and ON groups during either period. Optogenetic inhibition of c-Fos ensembles had no effect on tone-induced freezing. (Pre-tone: OFF vs ON, p=0.9850; Tone: OFF vs ON, p=0.8064, two-way repeated measures ANOVA with Sidak’s multiple comparisons test). (I) Quantification of freezing behavior during the context recall test. Optogenetic inhibition of c-Fos ensembles reduced contextual freezing (p = 0.0011, unpaired t-test). All data are shown as mean ± SEM. ns, not significant. *p< 0.05, **p<0.01. See also Table S1.

### Initial c-Fos ensembles in the DG become dispensable for memory retrieval after retraining

Previous studies suggest that repeated experiences can reorganize the neuronal populations involved in memory recall^16^. To test whether reorganization occurs within c-Fos ensembles in the DG following repeated training, FosTRAP mice received bilateral DG injection of AAV-EF1α-DIO-eNpHR3.0-EYFP to label c-Fos ensembles activated during initial training, which are hereafter referred to as ‘initial c-Fos ensembles’(Figure 2A). One day after the initial training, the mice underwent contextual fear conditioning under the same conditions (“Retraining”). Interestingly, optogenetic inhibition of the initial c-Fos ensembles did not affect the recall of memories enhanced by repeated training (Figures 2B and 2C), suggesting that memory enhancement through retraining may be supported by a neuronal population different from the initial c-Fos ensembles. We next performed a series of control experiments to test whether the initial c-Fos ensembles remained necessary when only specific components of the learning experience were repeated. Inhibiting these ensembles during recall led to reduced freezing when mice were re-exposed only to the context (Figures 2D and 2E) or received an immediate shock (Figures 2F and 2G), indicating that these manipulations did not induce ensemble reorganization. To determine whether any intervening learning event could alter the neuronal populations involved in recall, mice received a second training session in a novel context (Context C), distinct from the original one (Context A). We confirm that mice clearly discriminated between Context A and C (Figure S3A and S3B). Unlike retraining, this manipulation did not enhance freezing behavior (Figures S4A and S4B), and inhibition of the initial c-Fos ensembles also resulted in decreased freezing (Figures 2H and 2I). Therefore, when other events occurred between training and retrieval instead of retraining such as context re-exposure (Figures 2D and 2E), immediate shock (Figures 2F and 2G), or training in a distinct context (Figures 2H and 2I), the initial c-Fos ensembles were still required for memory recall, as observed in single training sessions (Figures 1C-1F). Together, these results suggest that while the initial DG c-Fos ensembles are required for memory retrieval under single-training or non-retraining control conditions, repeated contextual fear conditioning specifically drives the engram reorganization, rendering initial DG engram cells dispensable for retrieval.

**Figure 2.**
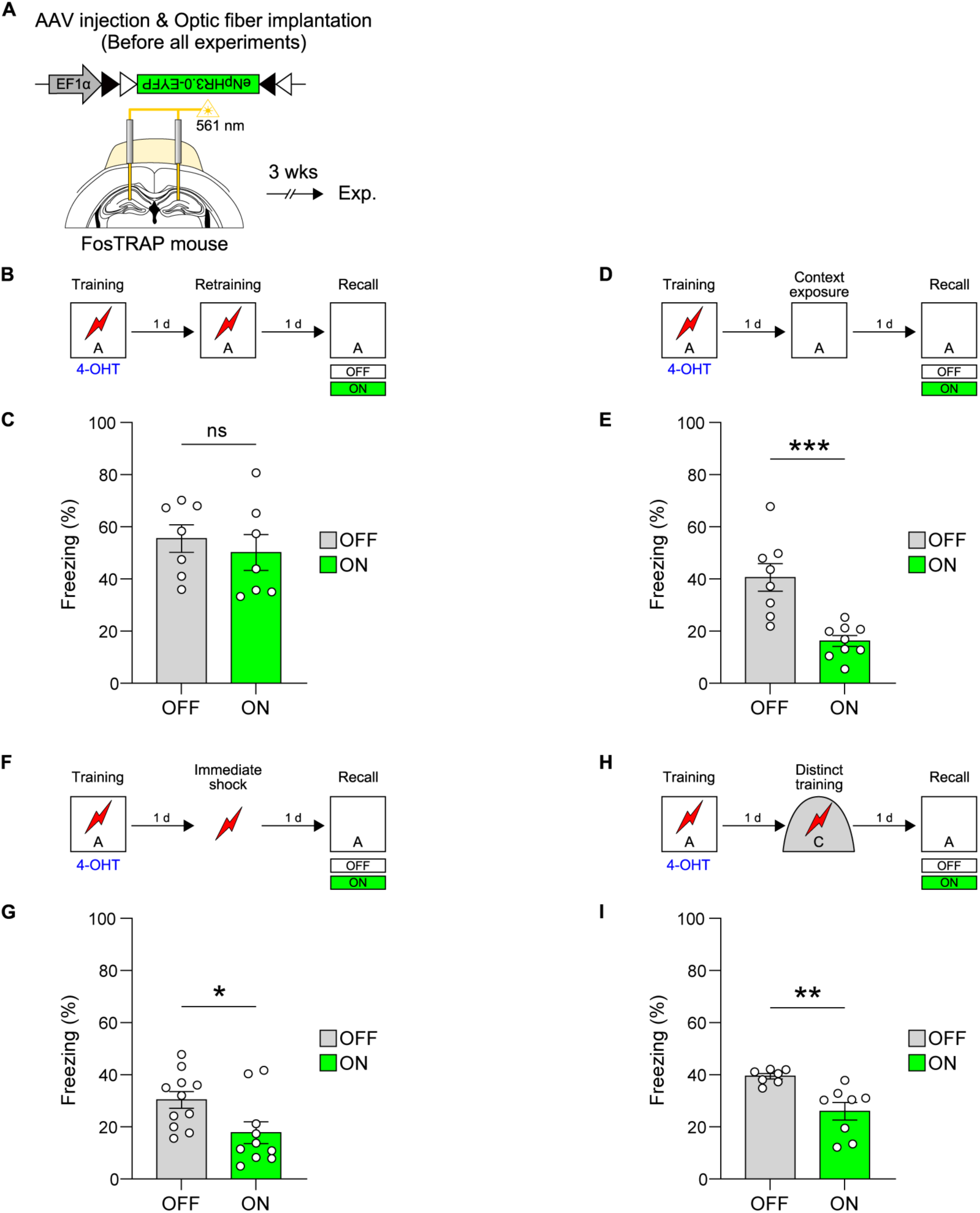
Initial c-Fos ensembles in the dentate gyrus become dispensable for memory retrieval after retraining (A) Schematic of surgery. AAV1-EF1α-DIO-eNpHR3.0-EYFP was bilaterally injected into the DG of FosTRAP mice, followed by implantation of optic fibers above the injection sites. Behavioral experiments were conducted three weeks later. (B) Behavioral procedure. Mice underwent training followed by 4-OHT injection to label initial c-Fos ensembles. After retraining, the ON group received light illumination during the context recall test, while the OFF group was tested without light (OFF group, n=7 mice; ON group, n=7 mice). (C) Freezing levels during recall were comparable between groups (p = 0.5504, unpaired t-test). (D) Behavioral procedure. Mice underwent training followed by 4-OHT injection to label c-Fos ensembles. Instead of retraining, mice were re-exposed to the context without shock. During the recall test, the ON group received light illumination, while the OFF group was tested without light (OFF group, n=8 mice; ON group, n=9 mice). (E) Optogenetic inhibition of initial c-Fos ensembles reduced freezing in the ON group compared to the OFF group (p=0.0004, unpaired t-test). (F) Behavioral procedure. Mice underwent training followed by 4-OHT injection to label c-Fos ensembles. Instead of retraining, mice received an immediate shock in the trained context. During the recall test, the ON group received light illumination, while the OFF group was tested without light (OFF group, n=11 mice; ON group, n=10 mice). (G) Optogenetic inhibition of initial c-Fos ensembles reduced freezing in the ON group compared to the OFF group (p=0.0197, unpaired t-test). (H) Behavioral procedure. Mice underwent initial training followed by 4-OHT injection to label c-Fos ensembles. A second training session was then conducted in a distinct context (Context C). During the recall test in the initial context, the ON group received light illumination, while the OFF group was tested without light (OFF group, n=7 mice; ON group, n=8 mice). (I) Optogenetic inhibition of initial c-Fos ensembles reduced freezing in the ON group compared to the OFF group (p=0.0034, unpaired t-test). All data are shown as mean ± SEM. ns, not significant. *p<0.05, **p<0.01, ***p<0.001. See also Table S1.

### Initial c-Fos ensembles in the DG display decreased spine densities after retraining

We next investigated whether synaptic remodeling underlies the reorganization of engram cell ensembles. To explore this idea, we quantified the dendritic spine density of initial DG c-Fos ensembles. To label the initial c-Fos ensembles, we used TRAP2; Ai14 mice (Figure 3A), in which activity-dependent neuronal labeling is achieved through tamoxifen-induced CreERT2-mediated recombination. TRAP2 mice^29^ express a tamoxifen-activated improved CreERT2, while Ai14 mice^34^ express tdTomato fluorescence upon Cre recombination. Following tamoxifen administration, neurons active during the labeling window were labeled with tdTomato (Figures 3B and 3C). Spine density was quantified in tdTomato+ (initial c-Fos ensemble) and tdTomato− neurons by injecting biocytin into individual neurons. As a control, mice receiving only a single training were included for the comparison. In the single training group, tdTomato+ neurons had significantly higher spine density than tdTomato− neurons (Figures 3D and 3E). In contrast, this training-induced increase in spine density was no longer observed in the retrained group (Figures 3F and 3G). This difference was not due to a time-dependent effect because spines were analyzed with the same delay after the initial training in both groups (Figure 3B). These findings suggest that synaptic-level changes in the initial c-Fos ensembles induced by the first training return to baseline levels following retraining, contributing to ensemble reorganization.

**Figure 3.**
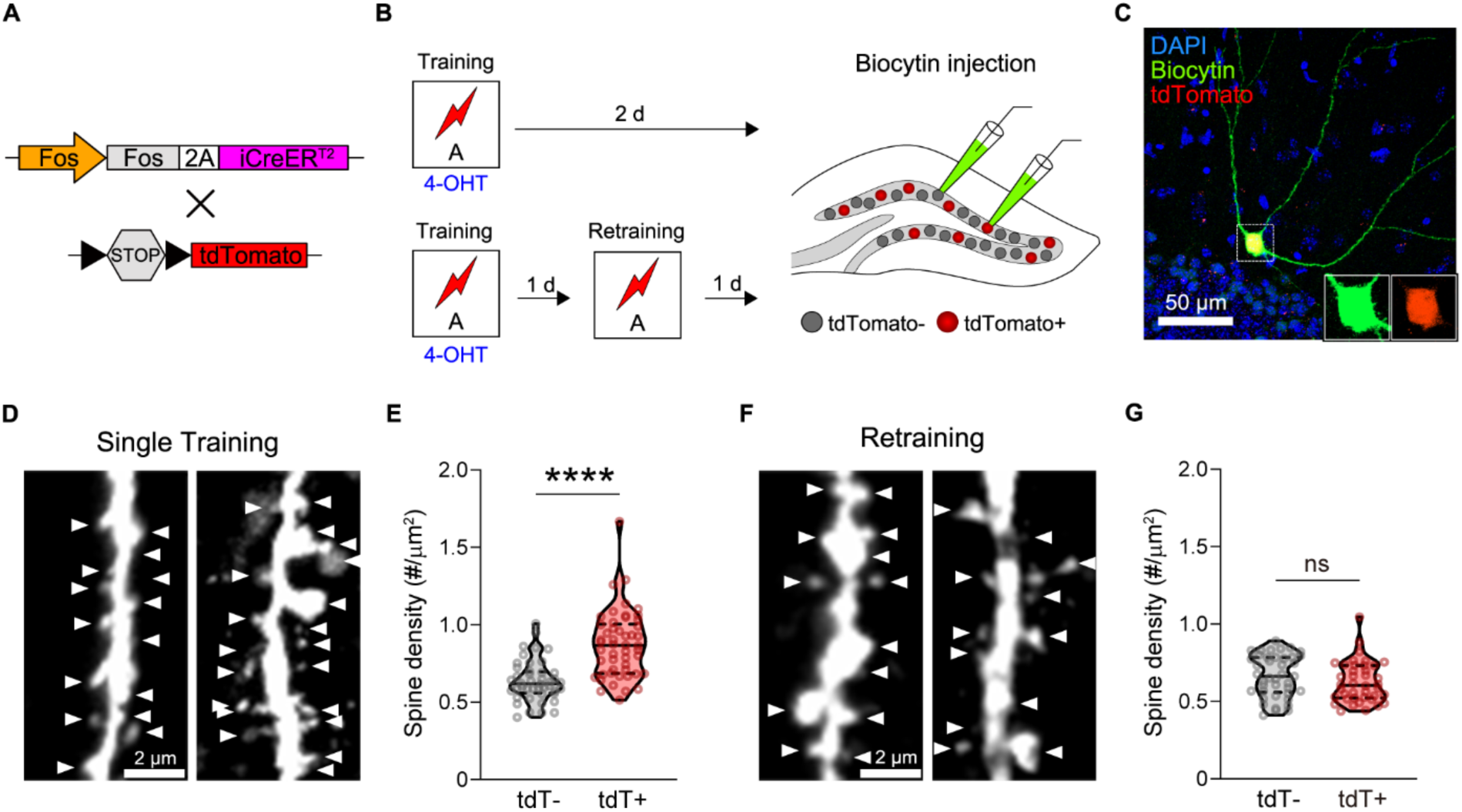
Loss of dendritic spines in initial DG engram cells by retraining (A) Schematic diagram of the TRAP2 x Ai14 mouse. Tamoxifen-induced Cre recombination in neurons active during a defined time window leads to permanent tdTomato expression. (B) Experimental procedure for spine analysis. The initial c-Fos ensemble was labeled by 4-OHT injection immediately after training. Mice in both the single training and retraining groups were sacrificed 2 days after labeling. Acute hippocampal slices were prepared, and tdTomato+ and tdTomato-DG neurons were identified under a fluorescence microscope. Individual neurons were patched and filled with biocytin for spine analysis. (C) Representative confocal image of DG neurons labeled during training (tdTomato+), filled with biocytin. Insets display magnified views of tdTomato+ neuron showing biocytin (green) and tdTomato (red) signals, respectively. Scale bar, 50 μm. (D and E) Spine analysis in the single-training group. (D) Representative confocal images of dendrites from tdTomato-(tdT-, left) and tdTomato+ (tdT+, right) DG neurons. Arrowheads denote the presence of dendritic spines. Scale bar, 2μm. (E) Average spine density in tdTomato- and tdTomato+ neurons (tdT-, n=44 cells; tdT+, n=50 cells; p<0.0001, Mann-Whitney test). (F and G) Spine analysis in the retraining group. (F) Representative confocal images of dendrites from tdTomato-(tdT-, left) and tdTomato+ (tdT+, right) DG neurons. Arrowheads denote the presence of dendritic spines. Scale bar, 2 μm. (G) Average spine density in tdTomato- and tdTomato+ neurons (tdT-, n=41 cells; tdT+, n=40 cells; p=0.1009, Mann-Whitney test). All data are shown as mean ± SEM. ns, not significant. ****p<0.0001. See also Table S1.

### Initial c-Fos ensembles in the DG are not preferentially reactivated during memory retrieval after retraining

We next determined whether reduced spine density is associated with decreased neuronal reactivation by using c-Fos expression as a proxy for neuronal activity during recall. Neurons activated during initial training were labeled with EYFP, and neurons active during retrieval were identified by c-Fos immunostaining (Figures 4A-4C). The size of EYFP+ labeled population reflected the number of neurons activated during the initial learning, and the c-Fos+ labeled population indicated the number of neurons activated during memory recall. Both the size of EYFP+ labeled population and c-Fos+ recall-activated population were similar between single training and retrained groups (Figure 4D and 4E). In addition, we confirmed no difference in the c-Fos expression levels induced by the single training and retraining (Figures S5A and S5B). Reactivation probability (the percentage of c-Fos+ cells among EYFP+ neurons) was significantly reduced in the retraining group compared with the single training group (Figure 4F). While the reactivation probability in the single training group was significantly above chance, that in the retraining group remained at chance level, indicating a lack of preferential reactivation of initial engram cells during memory recall. In the single-training group, the proportion of cells activated during retrieval (c-Fos+) revealed that EYFP+ neurons (cells active during training) showed a significantly higher activation rate than EYFP− neurons (cells not active during training) (Figure 4G). However, no such difference was observed between the two cell populations in the retraining group, indicating that initial c-Fos ensembles are less likely to be reactivated during memory retrieval after retraining (Figure 4H).

**Figure 4.**
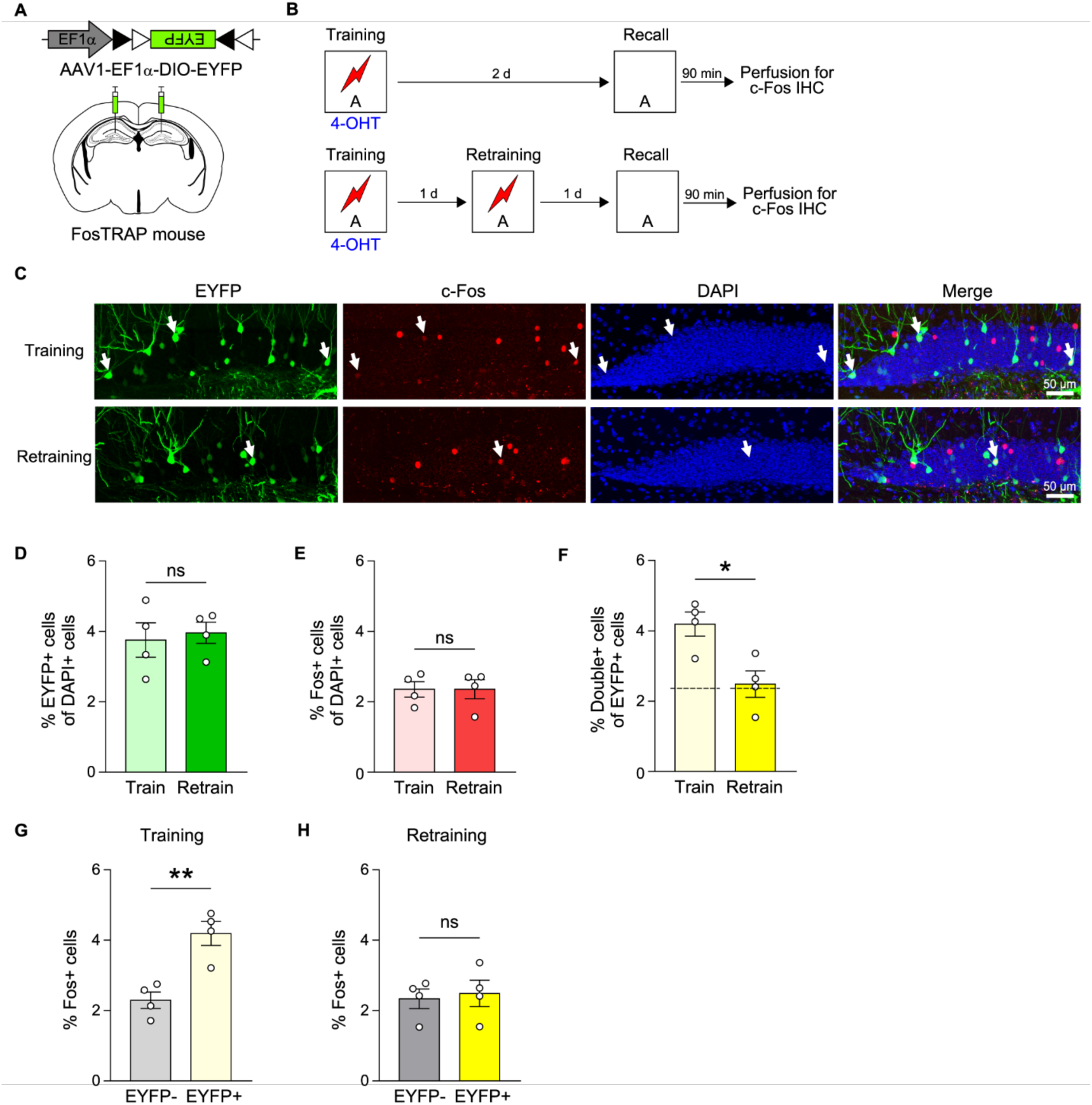
Reactivation of the initial DG engram cells decreases by retraining (A) Schematic of viral injection. AAV1-EF1α-DIO-EYFP virus injection into the bilateral DG. (B) Behavioral procedure for reactivation analysis in the DG. Mice in both the Training and Retraining groups were perfused 90 minutes after the recall session for c-Fos immunostaining (Training group, n=4 mice; Retraining group, n=4 mice). (C) Representative images of the DG. Confocal images from the Training (top) and Retraining (bottom) groups show EYFP+ neurons (green, activated during training), c-Fos+ neurons (red, activated during recall), and DAPI+ nuclei (blue). White arrows indicate double+ neurons. Scale bar, 50 μm (D) The size of initial c-Fos ensembles was comparable between the Training and Retraining groups (p=0.7332, unpaired t-test). (E) The percentage of neurons that were active during recall was comparable between the Training and Retraining groups (p>0.9999, unpaired t-test). (F) Quantification of reactivation probability. The proportion of double+ neurons among EYFP+ neurons was significantly higher in the Training group than in the Retraining group (p=0.0154, unpaired t-test). Dashed lines indicate the chance level for each group. Reactivation was significantly above chance in the Training group (p=0.0128, one-sample t-test), but not in the retraining group (p=0.7527, one-sample t-test). (G) In the Training group, c-Fos+ cell proportion was higher in EYFP+ than EYFP-neurons (p=0.0038, unpaired t-test) (H) In the Retraining group, no significant difference was observed in c-Fos+ cell proportion between EYFP+ and EYFP-populations (p=0.7550, unpaired t-test). All data are shown as mean ± SEM. ns, not significant. *p<0.05, **p<0.01. See also Table S1.

### c-Fos ensembles active during retraining in the DG are recruited for subsequent memory retrieval

Given that the initial ensemble becomes dispensable, we hypothesized that the memory is reorganized and stored in a distinct neuronal population recruited during the retraining. To test this hypothesis, we labeled and manipulated neurons active during the retraining. Unlike the previously presented experiments, the mice were administered 4-OHT immediately after retraining (Figures 5A and 5B). Mice receiving light illumination (ON group) displayed significantly reduced freezing behavior to the conditioned context compared to the control group with no light (Figure 5C). To determine whether this outcome was indeed caused by inhibiting neurons active at the time of retraining, we conducted control experiments. Mice were trained at the same time point as retraining, but in a distinct context, Context C (Figure 5D). In contrast to the retraining condition, inhibition of neurons activated during the second training in Context C did not result in memory impairment (Figure 5E). These results demonstrate that c-Fos ensembles activated during retraining are responsible for encoding repeated memories, suggesting a reshuffling of cell ensembles supporting memory in the DG.

**Figure 5.**
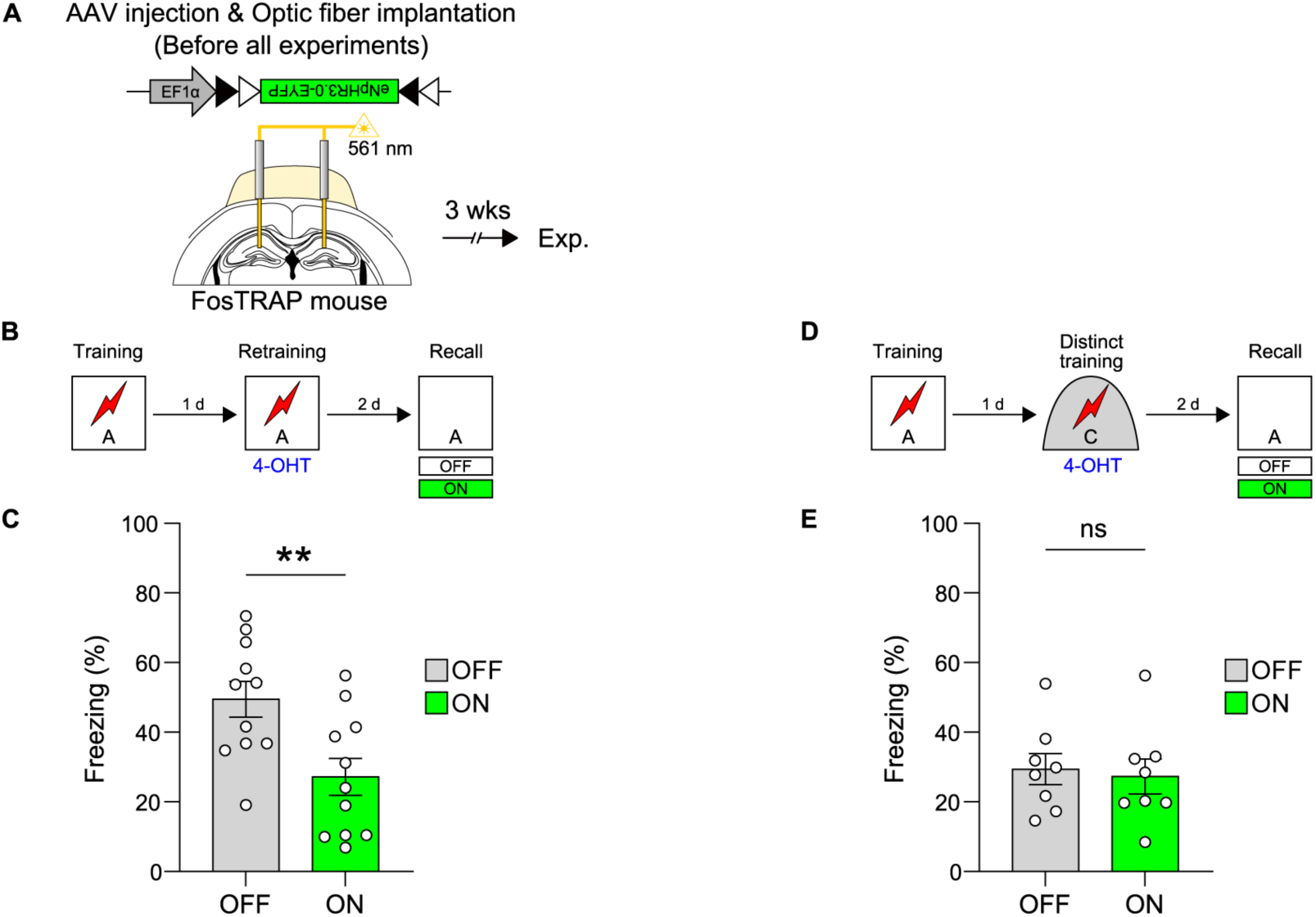
c-Fos ensembles activated during retraining in the DG are required for subsequent memory retrieval. (A) Schematic of surgery. AAV1-EF1α-DIO-eNpHR3.0-EYFP was bilaterally injected into the DG of FosTRAP mice, followed by implantation of optic fibers above the injection sites. Behavioral experiments were conducted three weeks later. (B) Behavioral procedure. c-Fos ensembles activated during retraining were labeled by 4-OHT injection immediately after retraining. During the context recall test, the ON group received light illumination, while the OFF group was tested without light (OFF group, n=11 mice; ON group, n=11 mice). (C) Optogenetic inhibition of retraining-activated ensembles reduced freezing in the ON group compared to the OFF group (p=0.0067, unpaired t-test). (D) Behavioral procedure. Mice were trained in a distinct context rather than retrained in the original context. c-Fos ensembles activated during the second training session were labeled by 4-OHT. During the context recall test, the ON group received light illumination, while the OFF group was tested without light (OFF group, n=8 mice; ON group, n=8 mice). (E) Freezing levels were comparable between OFF and ON groups (p=0.7572, unpaired t-test) All data are shown as mean ± SEM. ns., not significant. **p<0.01. See also Table S1.

### Reactivation of initial c-Fos ensembles in the mPFC during memory retrieval following retraining

Building on the well-established functional interaction between the hippocampus and the mPFC in episodic memory^35,36^, we next examined how retraining organizes engram cells in the mPFC. To this end, we labeled initial c-Fos ensembles in the mPFC (that is, cells activated during the initial training) and examined their reactivation during the retrieval test following retraining as in the hippocampus. We labeled the initial mPFC c-Fos ensembles with EYFP and assessed their reactivation during retrieval following retraining (Figures 6A and 6B). After retraining, mice were tested either in the conditioned context (Cond, recall in Context A) or the novel context (Novel, recall in Context B) (Figure 6B). The novel context group was included as a control for the non-memory retrieval condition. We confirmed that animals exhibited significant freezing specifically in the conditioned context compared to the novel context (Figure 6C). EYFP+ neurons marked those activated during the training, whereas c-Fos+ neurons identified cells activated during memory retrieval (Figures 6D and 6E). We then investigated how EYFP+ and EYFP-neurons were represented among activated cells during memory retrieval. There was no significant difference in the size of either EYFP+ or c-Fos+ populations between groups (Figures 6F and 6G). In the conditioned context group, EYFP+ neurons (cells activated during training; Figure 6H) showed a significantly higher reactivation probability compared to EYFP-neurons (cells not activated during training). In contrast, no such difference was observed between the two cell populations in the novel context group (Figure 6I). Therefore, following retraining, initial mPFC engram cells were selectively reactivated during memory retrieval.

**Figure 6.**
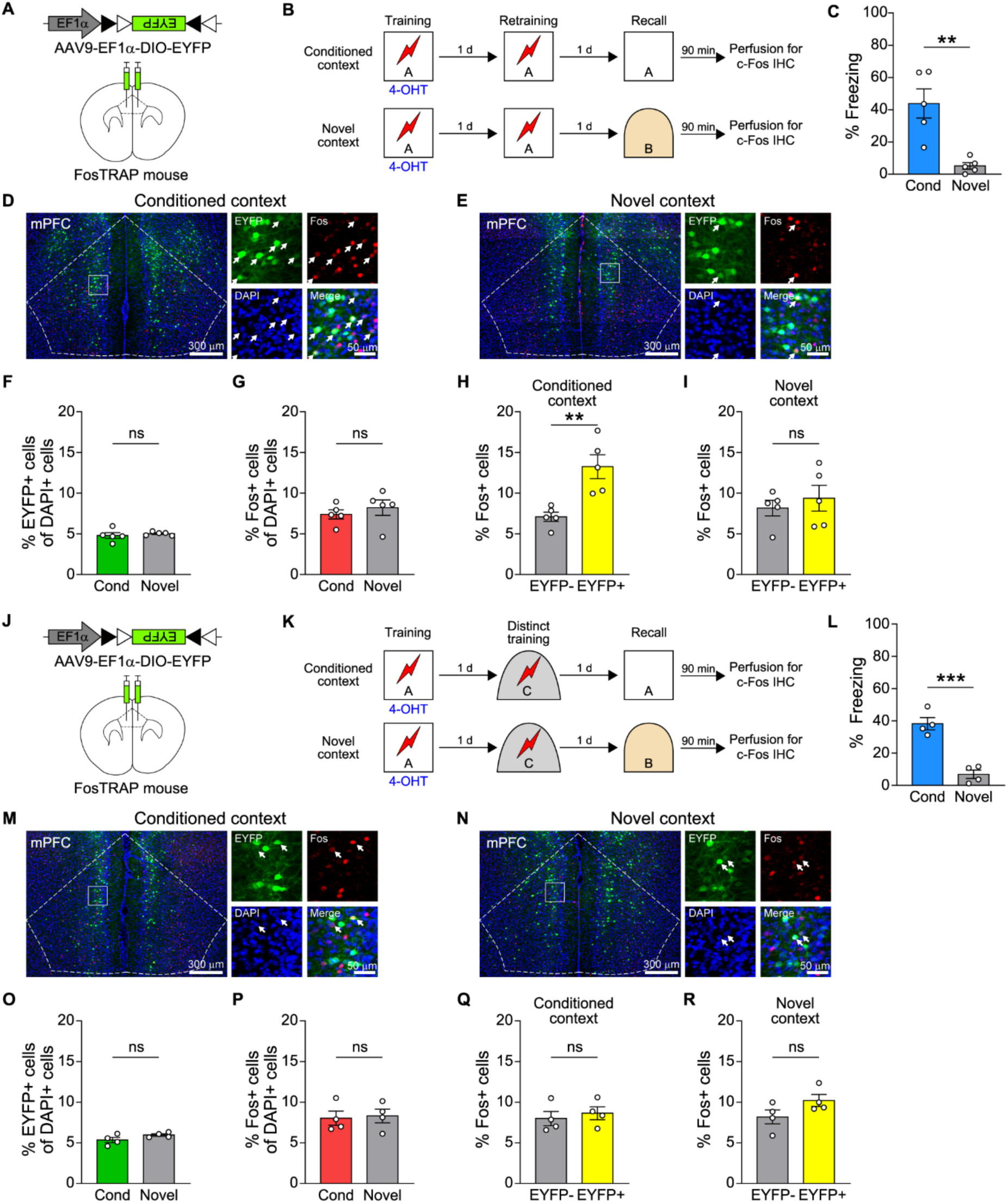
Initial mPFC engram cells are recruited during memory retrieval following retraining (A) Schematic of viral injection. AAV9-EF1α-DIO-EYFP was bilaterally injected into the mPFC. (B) Behavioral procedure for reactivation analysis in the mPFC after retraining. Neurons activated during the initial training were labeled with EYFP by 4-OHT injection. After retraining, mice were tested either in the conditioned context (Cond group, memory retrieval) or in a novel context (Novel group, no retrieval). Mice were perfused 90 minutes after the recall session for c-Fos immunostaining (Cond group, n=5; Novel group, n=5). (C) Freezing during the recall test was significantly higher in the Cond group than in the Novel group (p=0.0031, unpaired t-test) (D and E) Representative confocal images of the mPFC from Cond group (D), and Novel group (E). EYFP+ neurons (green, activated during training), c-Fos+ neurons (red, activated during recall), and DAPI+ nuclei (blue) are shown. Insets display magnified views with white arrows indicating double+ neurons. Scale bars, 300 μm (left) and 50 μm (insets, right). (F) The percentage of neurons that were active during recall was comparable between the Cond and Novel groups (p=0.4729, unpaired t-test). (G) The size of initial c-Fos ensembles was comparable between the Cond and Novel groups (p=0.5383, unpaired t-test). (H) In the Cond group, c-Fos+ cell proportion was higher in EYFP+ than EYFP-cell populations (p=0.0089, unpaired t-test) (I) In the Novel group, no significant difference was observed in c-Fos+ cells proportion between EYFP+ and EYFP-cell population (p=0.7444, unpaired t-test). (J) Schematic of viral injection. AAV9-EF1α-DIO-EYFP was bilaterally injected into the mPFC. (K) Behavioral procedure for reactivation analysis in the mPFC after second training in a distinct context. After the second training in a distinct context, mice were tested either in the conditioned context (Cond, memory retrieval group) or a novel context (Novel, no retrieval group). Both groups were perfused 90 minutes after the recall session for c-Fos immunostaining (Cond group, n=4; Novel group, n=4). (L) Freezing during the recall test was significantly higher in the Cond group than in the Novel group (p=0.0005, unpaired t-test). (M and N) Representative confocal images of the mPFC from Cond group (M), and Novel group (N). EYFP+ neurons (green, activated during training), c-Fos+ neurons (red, activated during recall), and DAPI+ nuclei (blue) are shown. Insets display magnified views with white arrows indicating double+ neurons. Scale bars, 300 μm (left) and 50 μm (insets, right). (O) The size of initial c-Fos ensembles was comparable between the Cond and Novel groups (p=0.1392, unpaired t-test). (P) The percentage of neurons that were active during recall was comparable between the Cond and Novel groups (p=0.8224, unpaired t-test). (Q and R) In both the Cond (Q) and Novel (R) groups, the proportion of c-Fos⁺ cells were comparable between EYFP+ and EYFP-cell populations (p=0.6011 and p=0.1240, respectively, unpaired t-test). All data are shown as mean ± SEM. ns, not significant. **p<0.01, ***p<0.001. See also Table S1.

To determine if this recruitment was specifically driven by retraining, we analyzed ensemble reactivation under a non-retraining condition in which mice were trained twice, as in the retraining paradigm, but in distinct contexts. FosTRAP mice received bilateral mPFC injection of AAV-EF1α-DIO-EYFP to label c-Fos ensembles active during initial training, and neurons that were active during recall were marked with c-Fos staining (Figures 6J and 6K). Following training in a distinct context (Context C), mice were assigned to one of two groups based on the context used for memory recall: conditioned context group (recall in Context A) or novel context group (recall in Context B) (Figure 6K). As shown above, we observed that animals exhibited significant freezing specifically in the conditioned context compared to the novel context (Figure 6L). There was no significant difference in the size of either the EYFP+ or c-Fos+ populations between groups (Figures 6M-6P). Notably, in contrast to what was observed following retraining, initial c-Fos ensembles were not preferentially reactivated during memory retrieval (Figures 6Q and 6R). These findings demonstrate that the recruitment of the mPFC engram cells is specifically driven by repeated learning.

### Rapidly recruited mPFC engram cells contribute to memory precision

Given that initial mPFC c-Fos ensembles were reactivated during retrieval following retraining, we examined whether these neurons play a functional role in memories updated by repeated learning. To this end, we inhibited these neurons during the memory retrieval test and assessed their effect on freezing. FosTRAP mice received bilateral mPFC injection of AAV-EF1α-DIO-EYFP or AAV-EF1α-DIO-eNpHR3.0-EYFP (Figures S6A-S6B). One day after retraining, the mice underwent a 6-minute context recall test in the conditioned context. During the first 3 minutes, we delivered a 561nm laser to inhibit the initial mPFC c-Fos ensembles (ON epoch), whereas no light was delivered during the last 3 minutes (OFF epoch) (Figure S6C). No significant differences were observed between ON and OFF epochs in either the EYFP or the NpHR group (Figures S6D and S6E). This indicates that inhibition of the initial mPFC c-Fos ensembles during memory recall had no significant effect.

Although inhibition of initial mPFC c-Fos ensembles did not alter freezing in the conditioned context, this does not necessarily mean that these neurons play no functional role in memory after retraining. We therefore asked whether the mPFC might instead contribute to memory precision. Importantly, whether repeated learning enhances memory precision or instead promotes fear generalization had remained unclear. To address this question, we tested B6J wild-type mice in two groups: a Training(Shock 2x) group, which received only two context-shock pairings without any retraining, and a Retraining group. One day after testing in the conditioned context (Context A), we assessed freezing in a highly similar context (Context A’) (Figures S7A and S7B). This context was identical to Context A except for the addition of a semicircular wall. In this condition, animals are required to discriminate subtle differences from the original environment and suppress inappropriate fear responses. The Retraining group showed a higher discrimination index than the Training (Shock 2x) group (Figure S7C). These results demonstrated that repeated learning improved the ability to discriminate the conditioned context from a highly similar one.

Based on this finding, we next performed an additional optogenetic inhibition experiment in both the conditioned context (Context A) and a similar context (Context A’). To inhibit initial mPFC c-Fos ensembles during the memory recall test, FosTRAP mice received bilateral mPFC injection of AAV-EF1α-DIO-EYFP or AAV-EF1α-DIO-eArch3.0-EYFP (Figures 7A and 7B). One day after retraining, mice underwent a 6-minute context recall test in the conditioned context, with the initial mPFC c-Fos ensembles inhibited during the first 3 minutes(ON epoch) and no light delivered during the last 3 minutes (OFF epoch) (Figure 7C). Freezing levels were comparable between the EYFP and Arch groups during either the ON or OFF epoch, and no significant differences were observed between ON and OFF epochs within each group (Figure 7D). These results are consistent with the experiment shown in Figure S6, where mPFC engram cells were inhibited using NpHR. One day after the recall test in the conditioned context, the same mice were tested in a highly similar context (Context A’), which was identical to Context A except for the addition of a semicircular wall. During the ON epoch, when the initial mPFC c-Fos ensembles were inhibited, the Arch group showed greater freezing than EYFP controls (Figure 7E). In contrast, no significant group differences were observed during the OFF epoch. Furthermore, within the Arch group, freezing levels significantly decreased from the ON to the OFF epoch, indicating that inhibition of the initial mPFC ensemble increased freezing specifically during light delivery. These results demonstrate that, after retraining, initial mPFC engram cells are crucial for maintaining memory precision.

**Figure 7.**
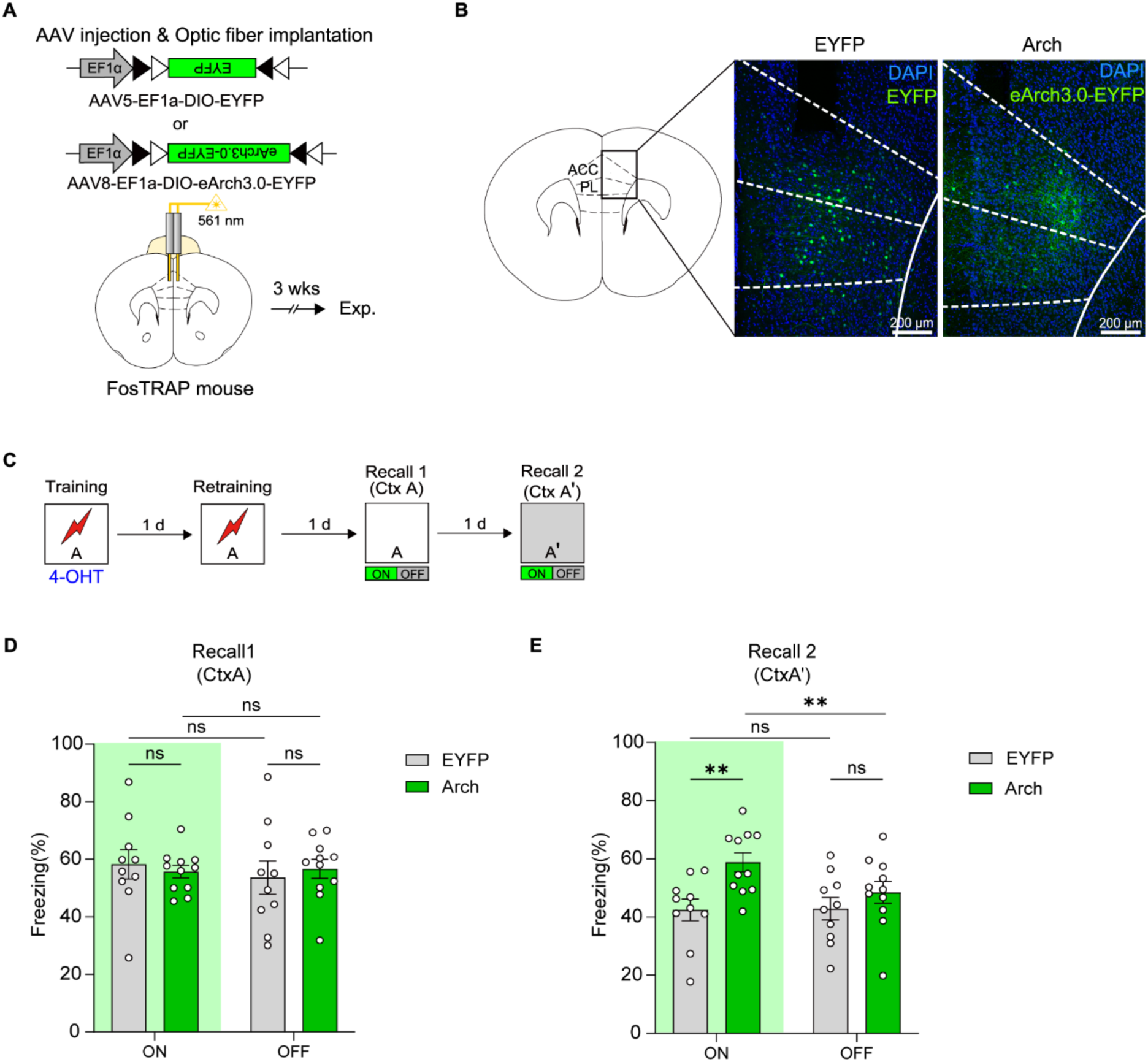
Initial mPFC c-Fos ensembles are required for memory precision in a similar context but not in the conditioned context (A) Schematic of surgery. AAV5-EF1α-DIO-EYFP or AAV8-EF1α-DIO-eArch3.0-EYFP was bilaterally injected into the mPFC, followed by implantation of optic fibers above the injection sites. Behavioral experiments were conducted three weeks after surgery. (B) Representative confocal images of the mPFC from EYFP group (left), and Arch group (right). Neurons activated during the initial training were labeled with EYFP or eArch3.0-EYFP by 4-OHT injection. Scale bars, 200 μm (C) Behavioral procedure for inhibition of initial mPFC c-Fos ensembles during conditioned and similar context recall tests. Mice underwent training followed by 4-OHT injection to label initial c-Fos ensembles. After retraining, mice were tested for memory recall in the conditioned context (Recall 1; Ctx A), followed by a recall test in a similar context (Recall 2; Ctx A’) on subsequent days. During both recall tests, light was delivered using an ON-OFF protocol, in which 561-nm laser light was applied during the first 3 min (ON epoch) and withheld during the subsequent 3 min (OFF epoch) (EYFP group n=10 mice, Arch group n=11 mice). (D) In the recall test of the conditioned context (Ctx A), freezing levels did not differ between EYFP and Arch groups during either the ON or OFF epoch. No significant within-group differences were observe between ON and OFF epochs (ON: EYFP vs Arch, p=0.8892; OFF: EYFP vs Arch p=0.8487; EYFP: ON vs OFF, p=0.1392; Arch: ON vs OFF, p=0.8963, two-way repeated measures ANOVA with Sidak’s multiple comparisons test). (E) In the similar context (Ctx A’), Arch group showed significantly higher freezing than EYFP group during ON epoch, whereas no difference was observed during the OFF epoch. Within-group analysis revealed no significant difference between the ON and OFF epochs in EYFP group, but a significant reduction in freezing from ON to OFF epoch in Arch group (ON: EYFP vs Arch, p=0.0062; OFF: EYFP vs Arch p=0.4875; EYFP: ON vs OFF p=0.9900; Arch: ON vs OFF p=0.0064, two-way repeated measures ANOVA with Sidak’s multiple comparisons test) All data are shown as mean ± SEM. ns, not significant. **p<0.01. See also Table S1.

### Reactivation of DG engram cells during relearning is critical for engram reorganization-dependent memory enhancement

Our results demonstrate that the engram cell population is reorganized in the hippocampal-mPFC network as associative learning is repeated. In contrast, it has been proposed that once recruited, the same engram cells are reused to encode the memory with repeated learning, based on the observation that inhibiting initially recruited engram cells disrupts memory enhancement induced by repeated training^23^. These seemingly inconsistent results may be explained if the reactivation of initial DG cells during relearning is critical for driving the reorganization, and that inhibiting them disrupts this reorganization process. This altered engram reorganization would then subsequently impair the retrieval of the enhanced memory, even if the acquisition of the second learning was successful.

To test this idea, we first examined whether reactivation of initial DG engram cells during retraining is necessary for retraining-dependent memory enhancement. We inhibited these neurons specifically during retraining and performed a recall test the following day. FosTRAP mice received bilateral DG injection of AAV-EF1α-DIO-eNpHR3.0-EYFP to label initial c-Fos ensembles (Figure 8A). Mice were assigned to either the Rt-OFF or Rt-ON group depending on whether they received 561nm laser illumination during the retraining session (Figure 8B). When freezing during the 3-minute pre-shock period of retraining was analyzed, the Rt-ON group showed reduced freezing compared to Rt-OFF group, confirming inhibition of the initial DG c-Fos ensembles (Figure 8C). This result is consistent with the data shown in Figure 1C and 1D. Notably, the Rt-ON group exhibited significantly reduced freezing compared to the Rt-OFF group (Figure 8D). Consistent with the previous report^23^, this result showed that initial DG c-Fos ensembles were required during retraining for memory enhancement induced by repetition.

**Figure 8.**
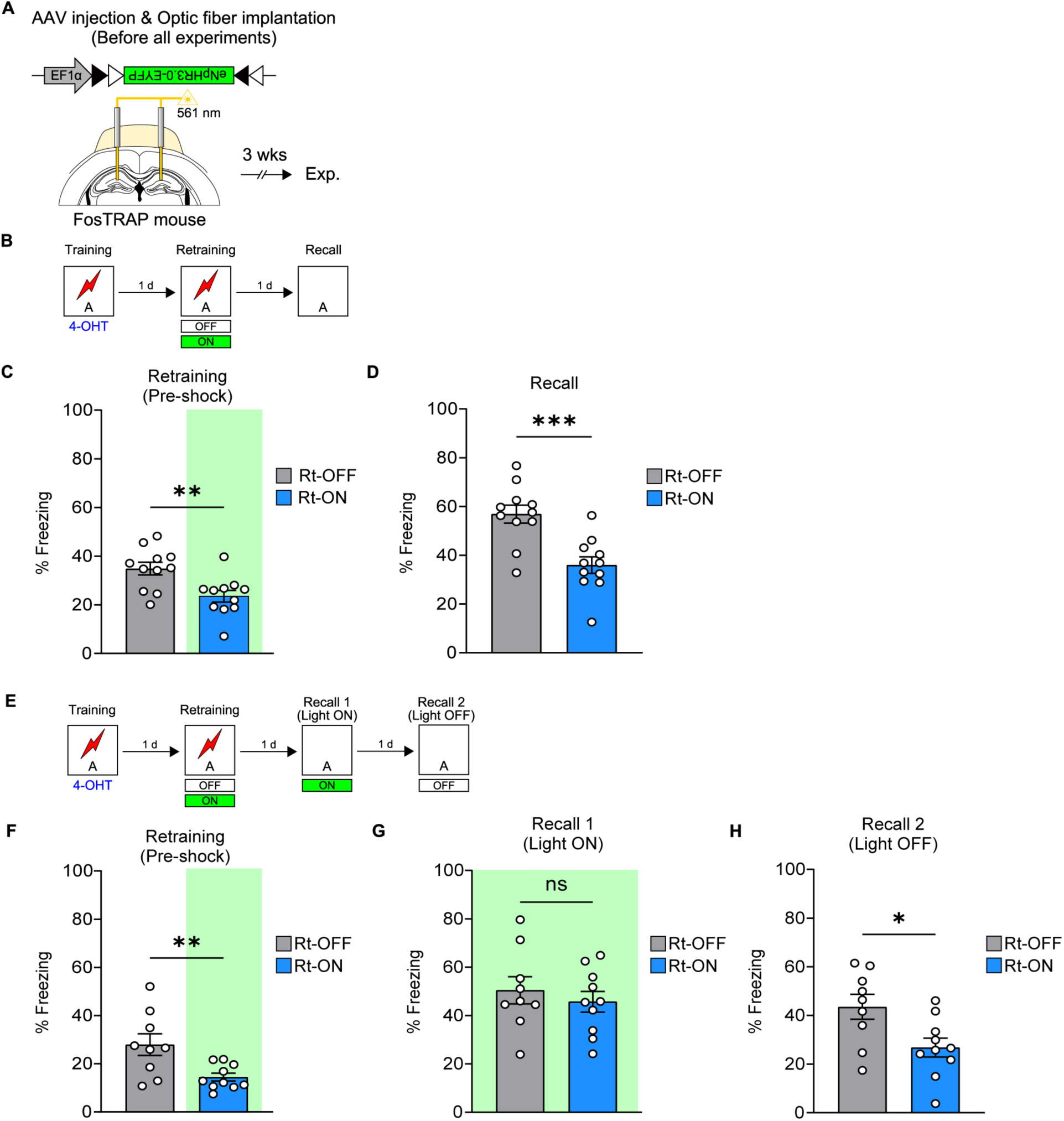
Reactivation of initial DG c-Fos ensembles during relearning is critical for engram reorganization-dependent memory enhancement (A) Schematic of surgery. AAV1-EF1α-DIO-eNpHR3.0-EYFP was bilaterally injected into the DG, followed by implantation of optic fibers above the injection sites. Behavioral experiments were conducted three weeks after surgery. (B) Behavioral procedure for inhibition of initial DG c-Fos ensembles during retraining. Mice underwent training followed by 4-OHT injection to label initial c-Fos ensembles. During retraining, the Rt-ON group received light illumination to inhibit the labeled neurons, whereas the Rt-OFF group was retrained without light stimulation. Context recall test was conducted the following day (Rt-OFF, n = 11; Rt-ON, n = 11). (C) Quantification of freezing behavior during retraining. Optogenetic inhibition of initial c-Fos ensembles reduced freezing during the pre-shock period of retraining in the Rt-ON group compared to the Rt-OFF group (p = 0.0047, unpaired t-test). (D) Quantification of freezing behavior during the context recall test. Rt-ON group showed significantly lower than Rt-OFF group during the context recall test (p = 0.0005, unpaired t-test). E) Behavioral procedure. Mice underwent training followed by 4-OHT injection to label initial c-Fos ensembles. Mice were divided into Rt-ON and Rt-OFF groups depending on whether they received light illumination during retraining. During retraining, the Rt-ON group received light stimulation to inhibit the labeled neurons, whereas the Rt-OFF group was retrained without light. Context recall tests were conducted on the following days: in Recall 1, all mice were tested under light illumination (Light ON); in Recall 2, conducted the next day, all mice were tested without light stimulation (Light OFF) (Rt-OFF, n = 9; Rt-ON, n = 10). (F) Quantification of freezing behavior during retraining. Optogenetic inhibition of initial c-Fos ensembles reduced freezing during the pre-shock period of retraining in the Rt-ON group compared to the Rt-OFF group (p = 0.0090, unpaired t-test). (G) In Recall 1 (all mice tested under 561 nm light illumination), freezing levels did not differ between the Rt-OFF and Rt-ON groups (p = 0.5046, unpaired t-test). (H) In Recall 2 (all mice tested without light illumination), freezing levels were significantly lower in the Rt-ON group than in the Rt-OFF group (p = 0.0170, unpaired t-test) All data are shown as mean ± SEM. ns, not significant. *p<0.05, **p<0.01, ***p<0.001. See also Table S1.

However, this finding also raises the possibility that inhibiting these neurons may have disrupted engram reorganization necessary for memory retrieval rather than preventing memory encoding for relearning. To distinguish these possibilities, we inhibited the same neurons during the recall test in Rt-ON and Rt-OFF groups (Figure 8E). Inhibition of the initial DG c-Fos ensembles was again confirmed by reduced freezing during the 3-minute pre-shock period (Figure 8F). Remarkably, when the initial DG c-Fos ensembles were inhibited again during recall following retraining, the Rt-ON group showed freezing levels comparable to those of the Rt-OFF group (Figure 8G), and this effect disappeared when the recall test was conducted the next day without light illumination (Figure 8H). These results indicate that even when the initial DG c-Fos ensembles were inhibited during retraining, new memory encoding for the retraining event still occurred but the initial DG engram cells interfered with the retrieval of transformed memory by relearning. Together, these findings support the idea that reactivation of the initial DG engram cells during relearning is critical for engram reorganization-dependent memory enhancement.

## DISCUSSION

Our findings identify a distinct system-wide engram reorganization driven by relearning, which differs from time-dependent reorganization. By directly manipulating these reorganized ensembles, our study provides the causal evidence linking this system-wide engram reorganization to the memory transformation, specifically the enhancement of memory strength and precision.

There have been inconsistent results about the role of the dentate gyrus in memory retrieval^11,27,30,37,38,39^. It appears that massive inhibition typically does not significantly impact memory retrieval, likely because compensatory routes are engaged and support memory retrieval. In contrast, inhibiting a sparse subset of neurons impairs performance. For instance, inhibiting Arc-induced DG neurons is shown to impair the retrieval of contextual fear memory^11^. Consistent with this result, we found that a sparsely labeled c-Fos neuronal ensemble is required for memory retrieval. Our findings thus reinforce the notion that the DG neurons inducing immediate-early genes (IEGs) by learning are engram cells required for later memory retrieval.

The reactivation of mPFC engram cells, which are typically involved in remote memory recall, during recent memory retrieval suggests an accelerated consolidation of cortical engram by repeated learning. A standard systems consolidation theory posits that memories initially depend on the hippocampal system but gradually transfer to the neocortex for long-term storage^40,41^. Previous studies in rodents and humans have suggested that memories can be rapidly consolidated into a form of cortex-dependent, hippocampus-independent memory, indicating accelerated systems consolidation^4,42,43,44,45^. Our results, however, cannot be explained by this process because we found active engram cells in both the hippocampus and the mPFC. To our knowledge, our study is the first to demonstrate the formation of dual engrams distributed across hippocampal-cortical networks by repeated experiences. Our findings align more closely with the multiple trace theory^46^, which posits that memory is quickly encoded in both the hippocampus and cortex in parallel, with both structures contributing to memory. Future studies should determine whether memories formed through repeated experiences exhibit different long-term dynamics or cortical dependencies compared to memories that simply become remote over time.

Although we observed reactivation of mPFC engram cells during memory retrieval, inhibiting them with optogenetics did not significantly affect freezing behavior during the retrieval test in the conditioned context. This is not surprising, however, given that the corresponding memory trace within the hippocampus remained intact and could compensate for cortical inhibition. Consistent with this interpretation, several previous studies have reported that suppression of mPFC or PL neurons tagged during learning does not impair memory retrieval at recent time points, suggesting that cortical engrams are not yet required for expression of recently formed memories^28,29,32^. While previous reports generally indicate that cortical engrams are dispensable at recent stages, a recent study demonstrated that prelimbic (PL) engram neurons activated during single contextual fear conditioning are already necessary for memory retrieval even at a recent time point, indicating that cortical engagement can occur earlier than previously thought^19^. In contrast, our repeated learning paradigm may induce a distinct form of systems reorganization in which the mPFC is rapidly engaged but assumes a different functional role. Rather than directly driving freezing behavior, mPFC engram neurons may contribute to the precision or discrimination of contextual memories that emerge through repeated experiences.

Indeed, when we inhibited mPFC engram cells during retrieval in a similar context, freezing behavior increased selectively in the Arch group, suggesting impaired discrimination between the conditioned and similar contexts. These findings suggest that cortical engagement through repeated learning confers new features on memory, such as precision, resulting in a repeat-dependent transformation of memory organization across hippocampal–cortical networks. By directly inhibiting the cortical ensembles that were recruited through repeated learning, our study reveals a functional contribution of these reorganized cells to a key feature of the transformed memory by repeated learning.

The detailed mechanism by which repeated learning reorganizes neural architectures to form dual engrams in hippocampal-cortical networks is unclear. Extensive research has documented the dynamic interaction between the hippocampus and the medial prefrontal cortex (mPFC) in memory formation^28,36,47,48,49^. These studies collectively indicate that simultaneous activation of both regions is essential not only for encoding new memories but also for consolidating remote ones. It has been suggested that increased post-learning offline reactivation in the cortex, as well as the coordinated offline reactivation between the hippocampus and cortex, may accelerate systems consolidation^4,44^. Moreover, artificially reactivating memory ensembles in the retrosplenial cortex (RSC) specifically during offline periods can accelerate systems consolidation^44^. Based on these findings, we hypothesize that increased offline reactivation of cortical c-Fos cells during sleep following repeated learning could drive a series of events that sculpt the reorganization of neural architectures supporting memory. It is unknown how the two distinct modes of engram reorganization in the hippocampus and mPFC (reshuffling and rapid maturation of engram cells, respectively) are coordinated. We speculate that silencing of the initial DG engram cells by the synapse loss or weakening may be critical for driving both changes. mGluR-LTD has been proposed as a mechanism for activity-dependent weakening of synapses in Arc-induced cells following repeated environmental exposure^50^. A similar mechanism may contribute to the silencing of the initial DG engram cells by repeated contextual conditioning.

Previously, we found that engram cells undergo turnover through repeated fear learning in the lateral amygdala^16^. Here, we demonstrate that the dentate gyrus also undergoes similar engram reorganization. However, this dynamic change is not a general feature of engram cells, as c-Fos cell ensembles in the mPFC remained stable. The difference in stability may reflect that the engram cells in the DG and mPFC play different roles for memory representations. DG engram cells may encode dynamic features of episodic events, while mPFC cells encode stable information, thereby contributing to context-specific memory retrieval. Although repetition, each event may have distinct episodic features. To encode the unique episodic features of each event, DG may create multiple traces by reorganizing engram cells.

Inhibition of initial DG engram cells during retraining impaired memory enhancement, indicating that their reactivation during retraining is critical for relearning. However, it was unclear whether encoding or retrieval was disrupted. We observed that memory retrieval following repeated learning was restored when these neurons were re-inhibited during recall. This suggests that other DG neurons were recruited to encode the retraining event even when the initial DG engram cells were inhibited, but that the initial DG engram cells interfered with the retrieval of transformed memory by relearning. Our findings thus suggest that reshuffling by silencing initial engram cells is critical for memory enhancement. How engram reorganization is altered by the inhibition of DG engram cells during retraining remains to be determined. We previously found that, unlike neuronal excitability-based memory allocation^51,52,53^, temporally distant repeated events (e.g., those with a 24-hour interval) appear to engage an excitability-independent memory allocation process^54^. The activity of existing engram cells may play a crucial role in guiding engram reorganization to support memory.

Taken together, our findings provide novel insights into how memory is distinctively shaped by time and experiences through a unique pattern of engram reorganization at the systems level, broadening our understanding of how engram cells support dynamic memory transformation and its relevance to neuropsychiatric disorders.

## RESOURCE AVAILABILITY

### Lead contact

Further information and requests for resources and reagents should be directed to the lead contact, Jin-Hee Han.

### Materials availability

This study did not generate new unique reagents.

## Data and code availability

All data and code that support findings of this study are available from the lead contact upon reasonable request.

## ACKNOWLEDGMENTS

We thank all members of the Memory Biology laboratory for helpful comments, discussions, and constructive suggestions. We thank the KAIST Bio-Core Center, the IBS center for Synaptic Brain Dysfunctions, and the KAIST Analysis Center for Research Advancement (KARA) for their assistance in performing confocal imaging experiments. M.K. was supported by the KAIST Jang Young Sil Fellow Program. This work was supported by the National Research Foundation of Korea (NRF) grant funded by the Korea government (MSIT) (RS-2023-NR077269).

## AUTHOR CONTRIBUTIONS

Conceptualization, M.K., B.S., and J.-H.H.; Methodology, M.K., B.S., Y.J., S.S., J.W.C., J.H., J.P., and J.-H.H.; Investigation, M.K., B.S., Y.J., S.S., J.W.C., J.H., and J.P.; Writing—original draft, B.S., M.K., and J.-H.H.; Writing—review & editing, B.S., M.K., and J.-H.H.; Funding acquisition, M.K. and J.-H.H.; Resources, J.-H.H.; Supervision, J.-H.H.

## SUPPLEMENTAL INFORMATION

Document S1. Figures S1–S7.

Table S1. Statistical details of all data related to STAR Methods.

## STAR★METHODS

### KEY RESOURCES TABLE

| REAGENT OR RESOURCE | SOURCE | IDENTIFIER |
| --- | --- | --- |
| Antibodies |  |  |
| Mouse anti-GAD67 | Millipore | Cat#MAB5406; RRID: AB_2278725 |
| Chicken anti-GFP | Aves | Cat#GFP-1010; RRID: AB_2307313 |
| Rabbit anti-c-Fos | Synaptic systems | Cat #226 003; RRID: AB_2231974 |
| Rabbit anti-c-Fos | Sigma | Cat# F7799; RRID: AB_259739 |
| Alexa Fluor 594-conjugated goat anti-rabbit | Invitrogen | Cat#A-11037; RRID: AB_2534095 |
| Alexa Fluor 488-conjugated goat anti-chicken | Invitrogen | Cat#A-11039; RRID: AB_142924 |
| Alexa Fluor 594-conjugated goat anti-mouse | Invitrogen | Cat#A-11005; RRID: AB_2534073 |
| Bacterial and virus strains |  |  |
| AAV1-EF1α-DIO-EYFP | Addgene | Item #27056-AAV1 |
| AAV5-EF1α-DIO-EYFP | Addgene | Item #27056-AAV5 |
| AAV9-EF1α-DIO-EYFP | Addgene | Item #27056-AAV9 |
| AAV1-EF1α-DIO-eNpHR3.0-EYFP | Addgene | Item #26971-AAV1 |
| pAAV-CaMKIIa-eArch3.0-EYFP | Addgene | Plasmid #35516 |
| AAV8- EF1α-DIO-eArch3.0-EYFP | N/A | N/A |
| Chemicals, peptides, and recombinant proteins |  |  |
| 4-hydroxytamoxifen | Sigma | Cat#H6278 |
| Experimental models: Organisms/strains |  |  |
| Mice: C57BL/6J (B6) | Jackson Laboratory | Strain #:000664; RRID:IMSR_JAX:000664 |
| Mice: FosTRAP (FosCreERT2) | Jackson Laboratory | Strain #:021882; RRID:IMSR_JAX:021882 |
| Mice: TRAP2 | Jackson Laboratory | Strain #:030323; RRID:IMSR_JAX:030323 |
| Mice: Ai14 | Jackson Laboratory | Strain #:007914; RRID:IMSR_JAX:007914 |
| Software and algorithms |  |  |
| FreezeFrame | Actimetrics | https://actimetrics.com/products/freezeFrame/ |
| ZEN lite software | Carl Zeiss | https://www.zeiss.com/microscopy/en/products/software/zeiss-zen-lite.html |
| Graphpad Prism 10 | GraphPad | https://www.graphpad.com/scientific-software/prism/ |
| IMARIS | Oxford instruments | https://imaris.oxinst.com/ |
| Fiji (ImageJ) | Schindelin et al. | https://imagej.net/software/fiji/downloads |

### EXPERIMENTAL MODEL AND STUDY PARTICIPANT DETAILS

#### Animals

C57BL/6J (JAX stock #000664), FosCreERT2 mice (JAX stock #021882), TRAP2 mice (JAX stock #030323) and Ai14 mice (JAX stock #007914) were obtained from the Jackson laboratory. To maintain a live mice colony, heterozygous mice were bred to wildtype C57BL/6J inbred mice. TRAP2 mice were bred with Ai14 mice to generate double heterozygous (TRAP2xAi14) mice, which were utilized in numerous experiments reported in a previous study^29^. Offspring heterozygous mice or homozygous mice were selected through genotyping. All male mice were used. In this study, FosTRAP mice primarily referred to FosCreERT2 mice. TRAP2 mice were used for spine analysis and Arch-mediated inhibition experiments, whereas all other experiments were conducted in FosCreERT2 mice. Mice aged 8 to 13 weeks were used in behavioral tests and were group-housed (3-5 mice per cage) until before stereotactic surgery. They maintained in 12 hours light/dark cycle at a constant temperature of 22 ± 2°C and 30-60% humidity. Food and water were provided ad libitum. Behavioral experiments were conducted during the light phase (7:00 A.M. to 7:00 P.M.) To minimize background tagging, mice were singly housed after virus injection and ferrule implantation. All procedures and protocols were approved by the KAIST Institutional Animal Care and Use Committee. All experiments were performed in accordance with the guidelines of the KAIST Institutional Animal Care and Use Committee.

## METHOD DETAILS

### Viruses

Adeno-associated virus (AAV) plasmid DNA used in this study was obtained from Addgene or generated by subcloning. AAV packaging was performed as previously described^55^. Briefly, HEK293T cells (ATCC #CRL-3216) were co-transfected with the AAV expression plasmid, helper plasmid pAdΔF6 and plasmid encoding AAV Rep/Cap genes (serotypes AAV2/1, AAV2/5, AAV2/8, or AAV2/9 using the calcium phosphate precipitation method. Following transfection, cells were incubated for 72 hours before harvest. AAV particles were purified using iodixanol gradient ultracentrifugation. Residual iodixanol was washed out using an Amicon centrifugal filter (Millipore, #UFC910024) with Dulbecco’s phosphate buffered saline (DPBS; Hyclon, #SH30264.01). Viral genome titers were quantified by quantitative polymerase chain reaction (QIAGEN, Rotor-Gene Q). Before virus injection, the viruses were diluted in 1X DPBS to a titer of 2–7× 10¹² vg/mL. The specific viral vectors and their final titers used in this study are as follows: 2 × 10¹² vg/mL for AAV1-EF1α -DIO-EYFP (Addgene, #27056-AAV1), 4 × 10¹² vg/mL for AAV5-EF1α -DIO-EYFP (Addgene, #27056-AAV5), 2 × 10¹² vg/mL for AAV9-EF1α -DIO-EYFP (Addgene, #27056-AAV9), 2.6 × 10¹² vg/mL for AAV1-EF1α -DIO-eNpHR3.0-EYFP (Addgene, #26966-AAV1), and 7 × 10¹² vg/mL for AAV1-EF1 α-DIO-eArch3.0-EYFP.

### Stereotactic surgery for virus injection and optic fiber implantation

Mice were anesthetized with pentobarbital (83 mg kg-1 of body weight) by intraperitoneal injection. Holes were drilled with an electrical driller at dorsal DG or mPFC. For labeling and manipulation of DG, virus was microinjected into two sites in the DG (injection site: From bregma, AP=-1.9mm; ML = ±1.25mm; DV - 1.95mm; 0.8μl/site). For labeling of mPFC, virus was microinjected into four sites (injection site 1&2: From bregma, AP=+1.7mm; ML = ±0.35mm; DV −1.8mm, injection site 3&4: From bregma, AP=+1.7mm ML = ±0.35mm; DV −2.2mm; 0.15μl/site). For manipulation of mPFC, virus was microinjected into two sites (From bregma, AP=+1.8mm; ML = ±0.35mm; DV −2.2 mm; 0.4μl/site). Virus solution was loaded into a glass pipette pre-filled with water and 3.0μl of mineral oil at the tip. After injection, the pipette was placed at the injection site for an additional 10 min to allow enough diffusion of the virus. Only for optogenetic inhibition experiments, mice were bilaterally implanted during the same surgery with optical fiber cannulas following virus injection. For dorsal DG targeting, bilateral mono optic fiber-optic cannulas (Doric lenses, 200 μm core diameter, 0.37NA, 2mm length) were implanted at the following stereotaxic coordinates from bregma: AP=-1.9mm; ML = ±1.25mm; DV −1.55mm. For medial prefrontal cortex(mPFC) targeting, dual optic fiber-optic cannula (Doric lenses, 200 μm core diameter, 0.37NA, length 2mm, pitch 0.7mm) were implanted at the following stereotaxic coordinates from bregma: AP=+1.8mm; ML = ±0.35mm; DV −1.9mm. As described previously^56^, to prevent visual disturbance caused by light leakage through the dental cement, optic fibers were secured using dental cement (Dentsply Sirona, #61118000) mixed with 20%(w/w) carbon powder (Sigma, #633100). After surgery, mice were placed on the heating pad until recovery and then returned to their home cages.

### Drug preparation

4-hydroxytamoxifen (4-OHT; Sigma, H6278) were dissolved through sonication in a solution of 10% ethanol and 90% corn oil (Sigma, C8267) to achieve a final concentration of 10 mg/mL. The preparation of 4-hydroxytamoxifen for TRAP2xAi14 mice or TRAP2 mice was carried out following a method adapted from DeNardo et al.^29^. To prepare the solution, 4-OHT was dissolved in ethanol at a concentration of 20 mg/ml by incubating at 37°C for 15 minutes with gentle shaking. A mixture of castor oil and sunflower seed oil (Sigma, Cat# 259853 and Cat# S5007, respectively) in a 1:4 ratio was then added to dilute the 4-OHT solution to a final concentration of 10 mg/ml. The ethanol was subsequently removed using vacuum centrifugation. The resulting 10 mg/ml 4-OHT solution was freshly prepared and used on the same day for intraperitoneal (i.p.) injections in all experimental procedures, administered at a dose of 50 mg/kg.

### Behavioral procedures

After 3–4 weeks of recovery following virus injection and fiber implantation, mice underwent a 6-day handling protocol for optogenetics experiments. During the first 3 days, animals were habituated to the experimenter to minimize stress, followed by 3 days of habituation to the fiber-optic cables. For experiments without optogenetic inhibition, mice underwent a 3-day handling protocol because fiber-optic cable habituation was unnecessary. This handling procedure was implemented to reduce experimental variability by alleviating anxiety in mice^57^.

Contextual fear conditioning was performed in a chamber with a grid floor connected to a shocker, all contained within a sound-attenuation box. We used five contexts for conditioning and recall tests. The alphabetical labeling of contexts (context A, A’, B, C) reflects their graded similarity to Context A. Context A consisted of a square chamber (Coulbourn Instruments, 18 x 18 x 30 cm) with a grid floor. Context A’ was located in a separate testing room and was identical to Context A except for the addition of semicircular walls. Context B was a context-shifted chamber (Coulbourn Instruments, 18 x 18 x 30 cm) with acrylic floor, semi-circular walls, vertically striped visual cues, beddings, and a 70% ethanol scent. Context C consisted of a larger square chamber (Med Associates, 30.5 × 24.1 × 21 cm) with a staggered-grid-floor, curved white plastic wall.

For the initial contextual fear conditioning session (Training), mice could explore the chamber freely for 3 minutes, after which a single foot shock (0.75 mA, 2 s) was delivered. Mice remained in the chamber for an additional 30 seconds before being removed and intraperitoneally injected with 4-hydroxytamoxifen (50 mg/kg) to tag neurons activated during conditioning. Following the injection, mice were returned to their home cages. Only during the single training (2x pairing) did mice receive two 2s 0.75mA foot shocks with 180-second inter-trial intervals.

A repeated contextual fear conditioning session (Retraining) was conducted 1 day after training, following the same protocol. For tagging neurons activated during retraining, mice were intraperitoneally injected with 4-hydroxytamoxifen (50 mg/kg) immediately after the retraining session. To evaluate memory recall, mice were returned to the conditioned chamber (Context A) or placed in a similar or novel chamber (Contexts A’, B, or C) for 3 minutes without receiving a foot shock. Freezing behavior during this period was measured to quantify recall of the contextual fear memory.

Optogenetic inhibition of the c-Fos ensembles was achieved by delivering 561 nm light upon the animal’s entrance into the chamber. Light delivery continued for the duration of the protocol. The laser intensity was calibrated to output 10-12mW (eArch3.0) or 12-15 mW (NpHR) at the tip of the optic fiber. Freezing behavior during the 3 minutes of exploration was recorded to assess the suppression of initial fear memory due to halorhodopsin (NpHR) inhibition. For mPFC experiments, the context test consisted of two consecutive 3-min sessions. Mice were placed in the designated context chamber, and freezing behavior was measured across both epochs. Continuous light was delivered through the optic fibers during the first 3-min session (ON epoch), whereas no light was applied during the subsequent 3-min session (OFF epoch).

### Auditory fear conditioning (AFC)

Auditory fear conditioning was performed in a chamber comprising a square cage (Coulbourn Instruments,18 × 18 × 30 cm) with a grid floor connected to a shocker, all contained within a sound-attenuation box. Following 3 min of exploration, the animals were exposed to a 30-second tone (white noise, 55dB) that ended simultaneously with a 2-second foot shock (0.75 mA). They remained in the chamber for an additional 30 seconds before being returned to their home cages. Two days later, mice were placed in context-shifted chamber with acrylic floor, semi-circular walls, vertically striped visual cues and 70% ethanol scent. The tone test consisted of a 3 min pre-tone baseline followed by a 90 s tone presentation. During the 90 s tone, the ON group received continuous 561 nm illumination via an optical fiber (200 µm core, NA 0.37; Doric Lenses), whereas the OFF group received no light. One day later, mice were returned to the original conditioning chamber for a 3 min context test. The ON group again received continuous 561 nm illumination throughout the session, while the OFF group received none.

### Analysis of Freezing

In optogenetic experiments, freezing behavior was manually scored due to the presence of optic fibers. The time mice remained immobile without any movement except breathing was counted as freezing behavior. Manual counting was performed in a blinded manner. The same manual scoring method was used in a previously published paper^58^. Percent freezing = time freezing/ total time x 100. In experiments that did not require optic fibers, freezing behavior was automatically scored using FreezeFrame software. Prism 10.4.2(GraphPad Software) was used for all statistical analyses in this study.

### Histology

To confirm viral expression and ferrule placement, mice were sacrificed following the completion of behavioral experiments, and brain sections were prepared. Anesthesia was induced with 2.5% avertin administered intraperitoneally, after which mice underwent transcardiac perfusion with phosphate-buffered saline (PBS) followed by cold 4% paraformaldehyde (PFA) for fixation. Post-perfusion, brains were immersed in 4% PFA overnight for additional fixation. Using a vibratome (VT-1200S, Leica Microsystems), coronal brain slices were sectioned at a thickness of 40 μm. These sections were mounted onto gelatin-coated slides with Vectashield antifade mounting medium (h-2000, Vector Laboratories). Images for verification purposes were acquired using a confocal microscope (LSM880 or LSM980, Carl ZEISS). Mice with off-target viral expression or mis-aligned ferrules, determined by referencing the Mouse Brain Atlas^59^, were excluded from the study.

### Spine analysis

To examine the structure of dendritic spines, we adopted an experimental approach based on the protocol outlined in Cho et al.^16^. A 0.1% biocytin solution was applied to EYFP-positive or EYFP-negative neurons in the dentate gyrus (DG) for 20 minutes. After biocytin labeling, brain slices were immersed in 4% paraformaldehyde (PFA) at 4°C for overnight fixation. The fixed slices underwent three washes with phosphate-buffered saline (PBS), followed by a 1-hour incubation at room temperature in a blocking solution comprising 5% goat serum and 0.2% Triton X-100 in PBS. Subsequently, the slices were treated with a streptavidin-AlexaFluor-594 conjugate (1:1000, Invitrogen) overnight at room temperature. After three additional washes in PBS, the slices were mounted on gelatin-coated slides using a DAPI-based mounting medium (h-1200, Vector). Imaging focused on secondary or tertiary dendritic branches located 50–100 μm from the soma of pyramidal neurons within the DG. Z-stack images were captured at 0.2 μm intervals using a Zeiss confocal microscope (models LSM780 or LSM880) equipped with a 63X objective lens, available through KAIST’s Bio-Core and Analysis Centers. Spine density measurements were performed using ImageJ software on 10 μm segments of dendrites.

### Immunohistochemistry

For GAD67 immunostaining, we used a detergent-free protocol as previously described^60^. Brain sections were first permeabilized for 30 minutes in 0.1% Triton X-100, then blocked in detergent-free blocking buffer (5% goat serum, 3% BSA in PBS). Sections were incubated with mouse anti-GAD67 primary antibody (MAB5406, Millipore; 1:500) at room temperature for 48 hours. After four 10 min of PBS washing, sections were incubated with Alexa Fluor 594–conjugated goat anti-mouse secondary antibody (1:2000) for 2 hours at room temperature. Following four additional 10-min washes in PBS, brain sections were mounted on gelatin-coated slides and counterstained with DAPI mounting medium (h-2000, Vector).

For c-Fos staining, mice were perfused 90 min following the context test (Recall). Brains were extracted and stored in 4% PFA overnight. Coronal brain sections (40μm) were obtained using a vibratome (VT-1200S, Leica). eNpHR3.0-EYFP signal was amplified with immunostaining. Brain sections were washed in PBS three times and were blocked for 1 hour at RT in blocking buffer containing goat serum. Sections were incubated with primary antibodies (1:2000 rabbit anti-c-Fos [226 003, Synaptic systems]; 1:5000 chicken anti-GFP [GFP-1020, Aves labs]) and allowed to incubate on a shaker overnight at room temperature. After four 10 min of PBS washing, sections were then incubated with secondary antibodies (1:2000 Alexa Fluor 594 anti-rabbit [Invitrogen]; 1:2000 Alexa Fluor 488 anti-chicken [Invitrogen] or 1:2000 Alexa Fluor 488 anti-rabbit [Invitrogen]) for 2 hours at RT. Following four additional 10-min washes in PBS, brain sections were mounted on gelatin-coated slides and counterstained with DAPI mounting medium (h-2000, Vector).

For DAB (diaminobenzidine) staining, brains were obtained 90 min after training or retraining. Anti-c-Fos primary antibodies (1:5000; F7799, Sigma) were used to detect c-Fos, which was followed by incubation with biotinylated goat-anti-rabbit secondary antibody (1:2000; Vector laboratories). After applying avidin-biotin based peroxidase system coupled with DAB (D5905, Sigma) as a chromogen, sections were mounted on the slides, dehydrated, and covered with cytoseal (8311-4, Thermo Scientific).

### Cell counting analysis

Based on the Mouse Brain Atlas^59^, the DG was defined in coronal sections from AP –1.7 mm to –2.3 mm, and the mPFC from AP +1.4 mm to +1.9 mm. For each animal, we analyzed six DG sections and five mPFC sections, averaging the percentage of labeled cells across those sections. Confocal microscopy was used to acquire z-stack images of the selected sections. DG Cell counting was performed using IMARIS 8 software (Bitplane). DAPI+ cells were identified using the spot detection algorithm in the IMARIS software. In contrast, the total number of EYFP+ cells, c-Fos+ cells, DAPI+ cells, and double-labeled cells (EYFP+ and c-Fos+) were manually counted to ensure accuracy. DG cell counting following DAB staining and mPFC reactivation analysis were performed using ImageJ.

## QUANTIFICATION AND STATISTICAL ANALYSIS

All statistical analyses in this study were performed using GraphPad Prism version 10.4.2. The Shapiro-Wilk test was used to assess the normality of data distribution. Depending on the results of the normality assessment, comparisons between two groups were conducted using either parametric tests (two-tailed unpaired test) or non-parametric tests (Mann–Whitney test). For analyses involving more than two conditions, one-way ANOVA or two-way repeated-measures ANOVA was applied. The latter was used when two groups were evaluated across repeated within-subject conditions. When ANOVA indicated significant effects, post-hoc pairwise comparisons were performed using Tukey’s or Sidak’s multiple-comparison procedures, as appropriate. Statistical significance was determined at a threshold of p < 0.05.

