## Supplemental data for "Repeated experience drives multiscale engram reorganization to shape memory strength and precision"

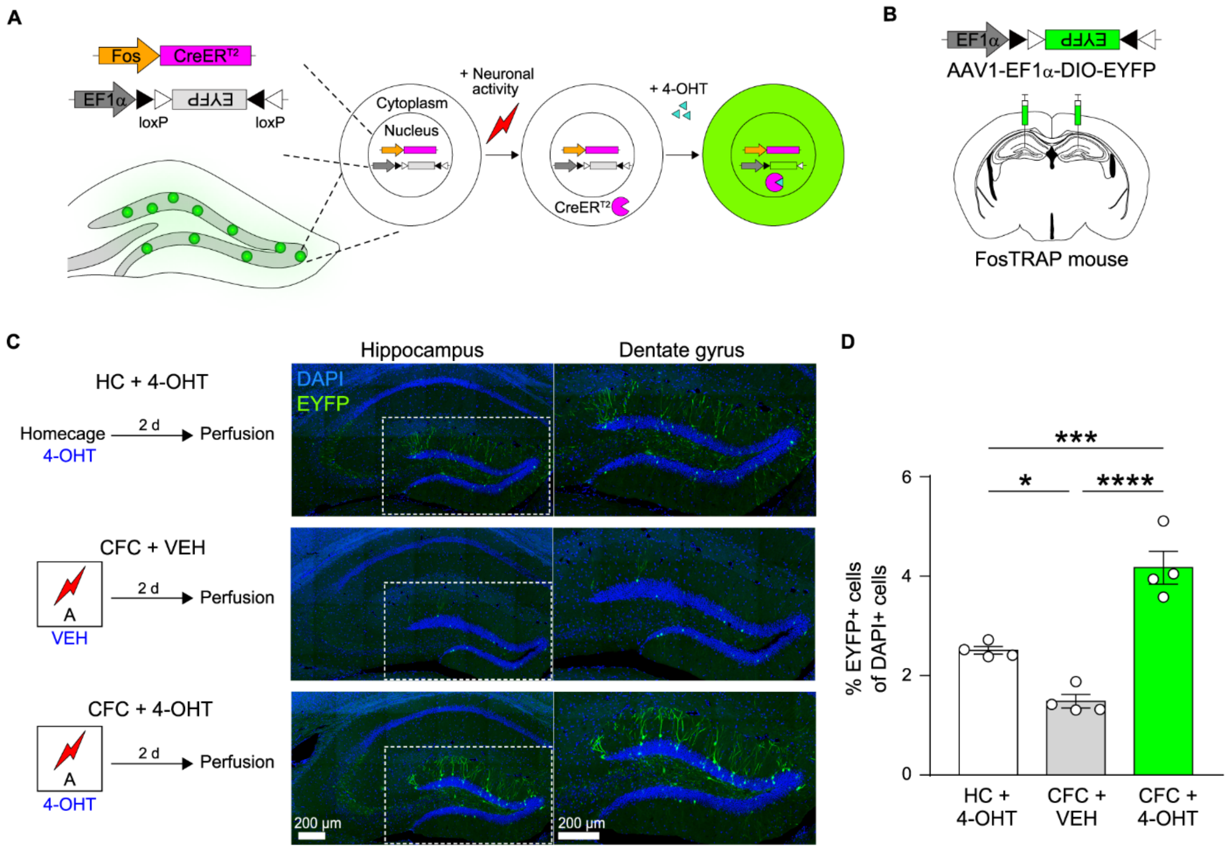


**Figure S1. c-Fos ensembles labeling in the DG**

(A) Schematic of a 4-OHT(4-hydroxytamoxifen)-driven strategy using an AAV expression vector to selectively label activated (Fos+) cells in FosTRAP (FosCreERT2) mice

(B) AAV1-EF1α-DIO-EYFP virus injection into the DG. Three weeks after virus injection, behavioral experiment progressed.

(C) Representative images in three groups: (top)HC+4-OHT (animals that remained in their home cages before receiving 4-OHT, n=4), (middle) CFC+ VEH (animals that received vehicle injections after CFC, n=4), and (bottom) CFC+ 4-OHT (animals that received 4-OHT immediately after CFC, n=4). The scale bar indicates 200μm.

(D) Quantification of EYFP-positive DG cells (EYFP+ of DAPI+ percentage) in three groups. Compared to the other two groups (HC+4-OHT, CFC+VEH), CFC+4-OHT group exhibited a significantly higher number of EYFP-positive cells. (HC+4-OHT vs CFC+VEH $p=0.0180$, HC+4-OHT vs CFC+4-OHT $p=0.0009$, CFC+VEH vs CFC+4-OHT $p<0.001$)

Data are shown as mean ± SEM. *$p < 0.05$, ***$p < 0.001$, ****$p < 0.0001$ by One-way ANOVA with Tukey's multiple comparisons test. See also Table S1.

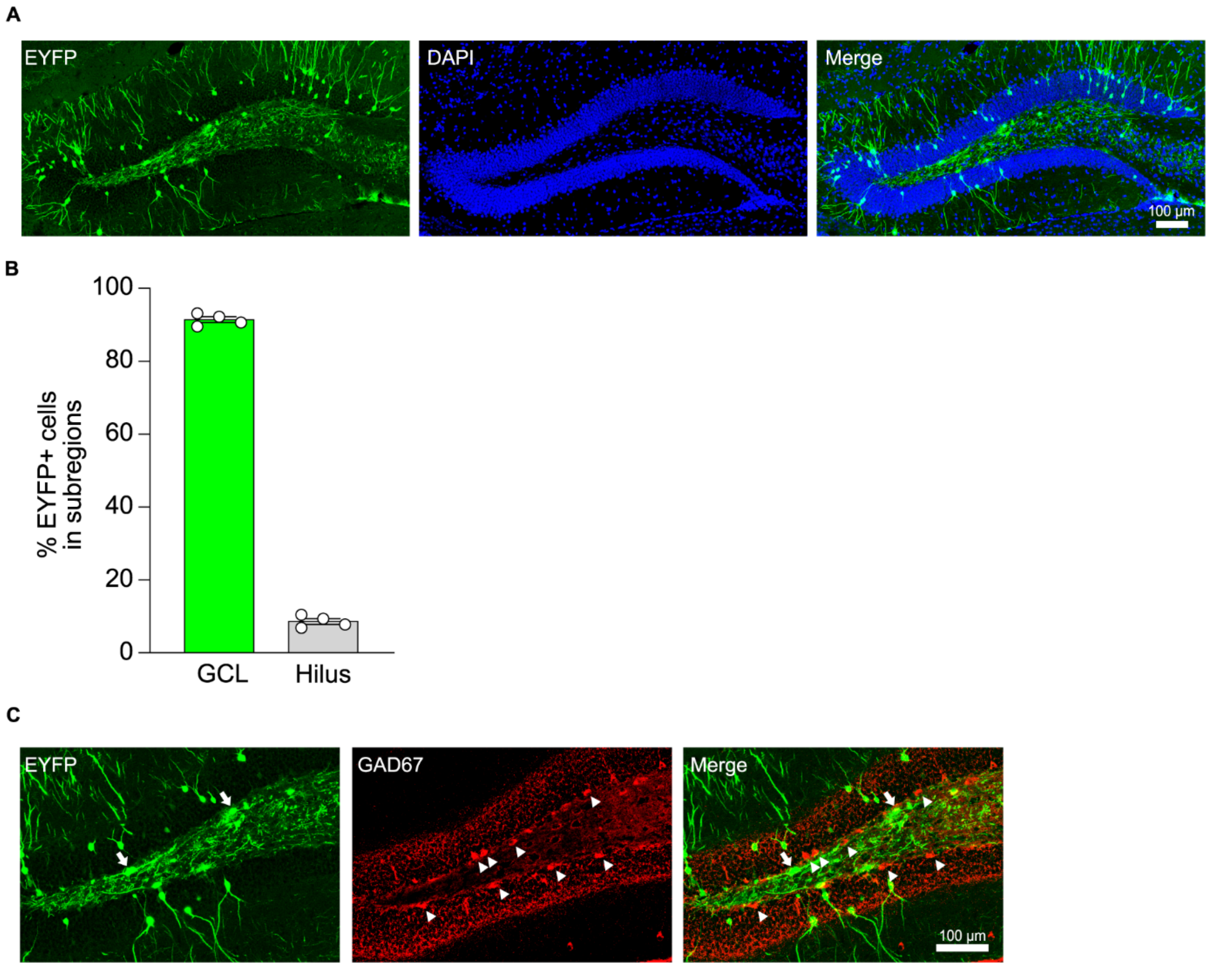


**Figure S2. 4-OHT administration and learning-dependent labeled neurons consist of excitatory neurons in the granule cell layer**

(A)Representative images of EYFP (left), DAPI (middle), and merged image (right) from CFC+ 4-OHT (animals that received 4-OHT immediately after CFC, n=4). The scale bar indicates 100µm.

(B) Quantification of the percentage of EYFP-positive cells in the granule cell layer (GCL) and hilus.

(C) Representative images of EYFP-positive cells (left; white arrows), GAD67-positive cells (middle; arrowheads) and merged image(right) showing no colocalization. The scale bar indicates 100µm.

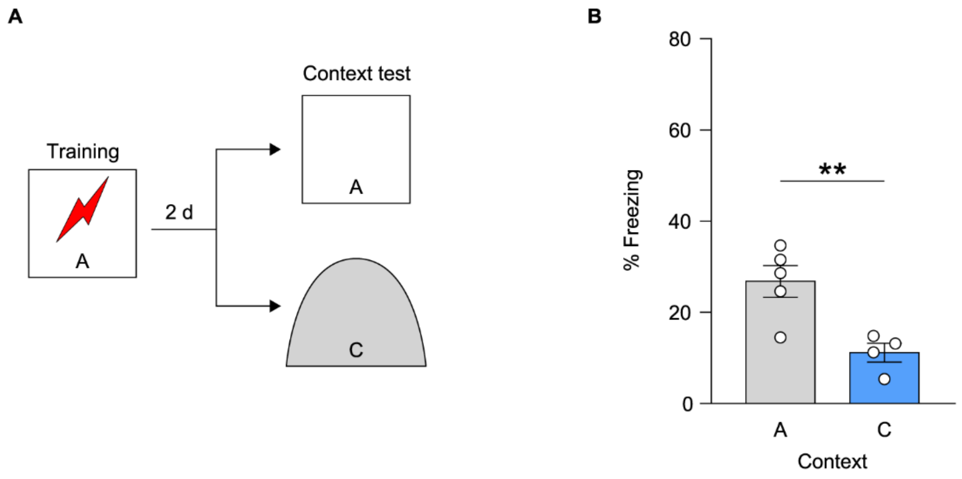


**Figure S3. Context discrimination in context A and C**

(A) Behavioral experiment scheme for context discrimination. Two days after training, mice were assigned to either the Context A or Context C test group depending on the context in which the recall test was performed.

(B) Freezing was measured during context recall test. (Context A n=5, Context C n=4; p=0.0088)

Data are shown as mean ± SEM. **p < 0.01 by Two-tailed unpaired t-test. See also Table S1.

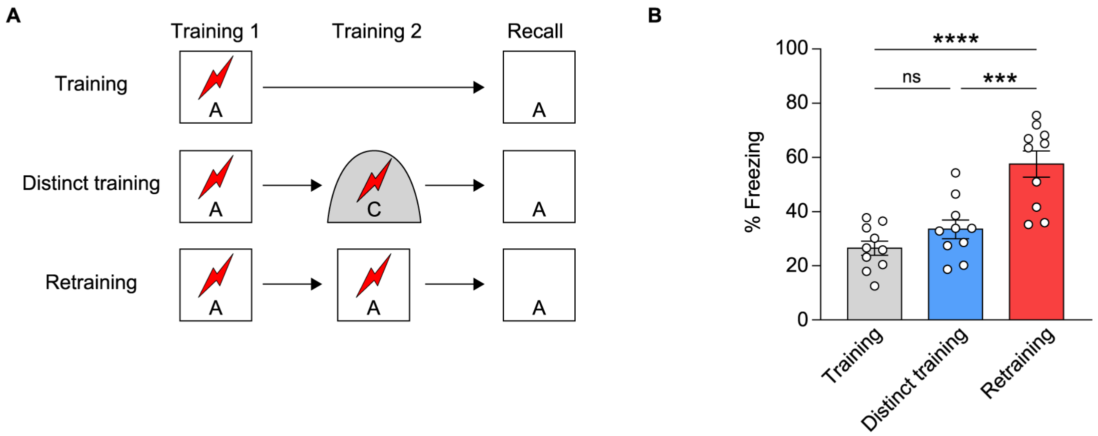


**Figure S4. Retraining enhances memory recall, but not second conditioning in distinct context does not**

(A) Behavioral experiment scheme for the Training, Distinct Training and Retraining groups

(B) Freezing was measured during context recall test (Training (Tr) n=10, Distinct training (Dt) n=10, Retraining (Rt) n=10; Tr vs Dt p=0.3969, Tr vs Rt p<0.0001, Dt vs Rt p=0.0003)

All data are shown as mean ± SEM. n.s., not significant. ***p < 0.001 and ****p < 0.0001 by One-way ANOVA with Tukey's multiple comparisons test. See also Table S1.

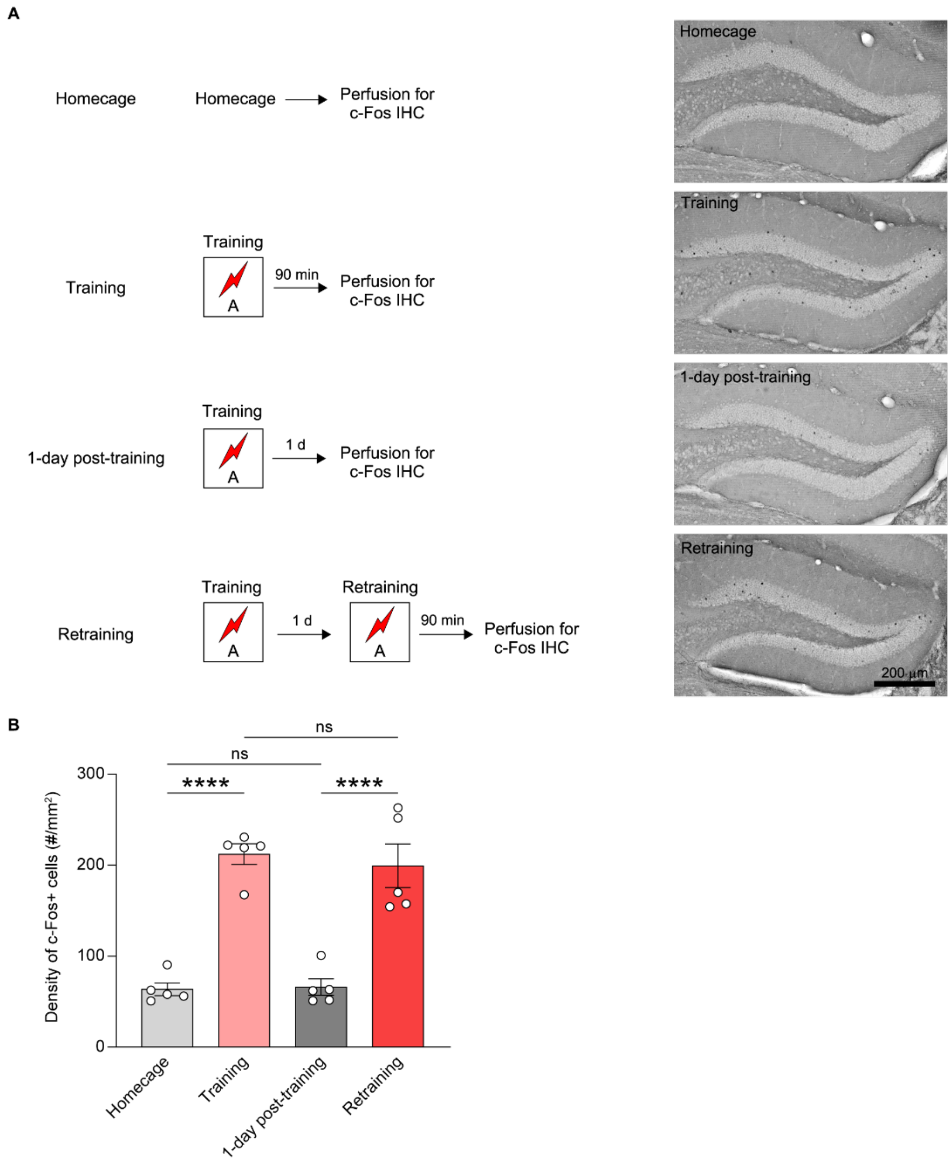


**Figure S5. Comparable levels of Fos positive cells are activated during contextual fear conditioning and re-conditioning in the DG**

(A) Representative bright filed microscope images show cFos expression (visualized by DAB staining) in the DG across four groups: (top) HC (animals that remained in their home cages and were perfused 90 minutes later. (upper-middle) Training (animals that underwent contextual fear conditioning once and were perfused 90 minutes later. (lower-middle) 1-day post-training (animals that underwent contextual fear conditioning once and were perfused 1 day later. (bottom) Retraining (animals that underwent re-conditioning and were perfused 90 minutes later). Scale bar, 200μm.

(B) Density of Fos+ cells (#/mm2) in the DG (HC n=5, Training n=5, 1-day post-training n=5, Retraining n=5; HC vs Training $p<0.0001$, HC vs 1 day post-training $p=0.9995$, HC vs Retraining $p<0.0001$, Training vs 1 day post-training $p<0.0001$, Training vs Retraining $p=0.9208$, 1 day post-training vs Retraining $p<0.0001$).

Data are shown as mean ± SEM. n.s., not significant. ****$p < 0.0001$ by One-way ANOVA with Tukey's multiple comparisons test. See also Table S1.

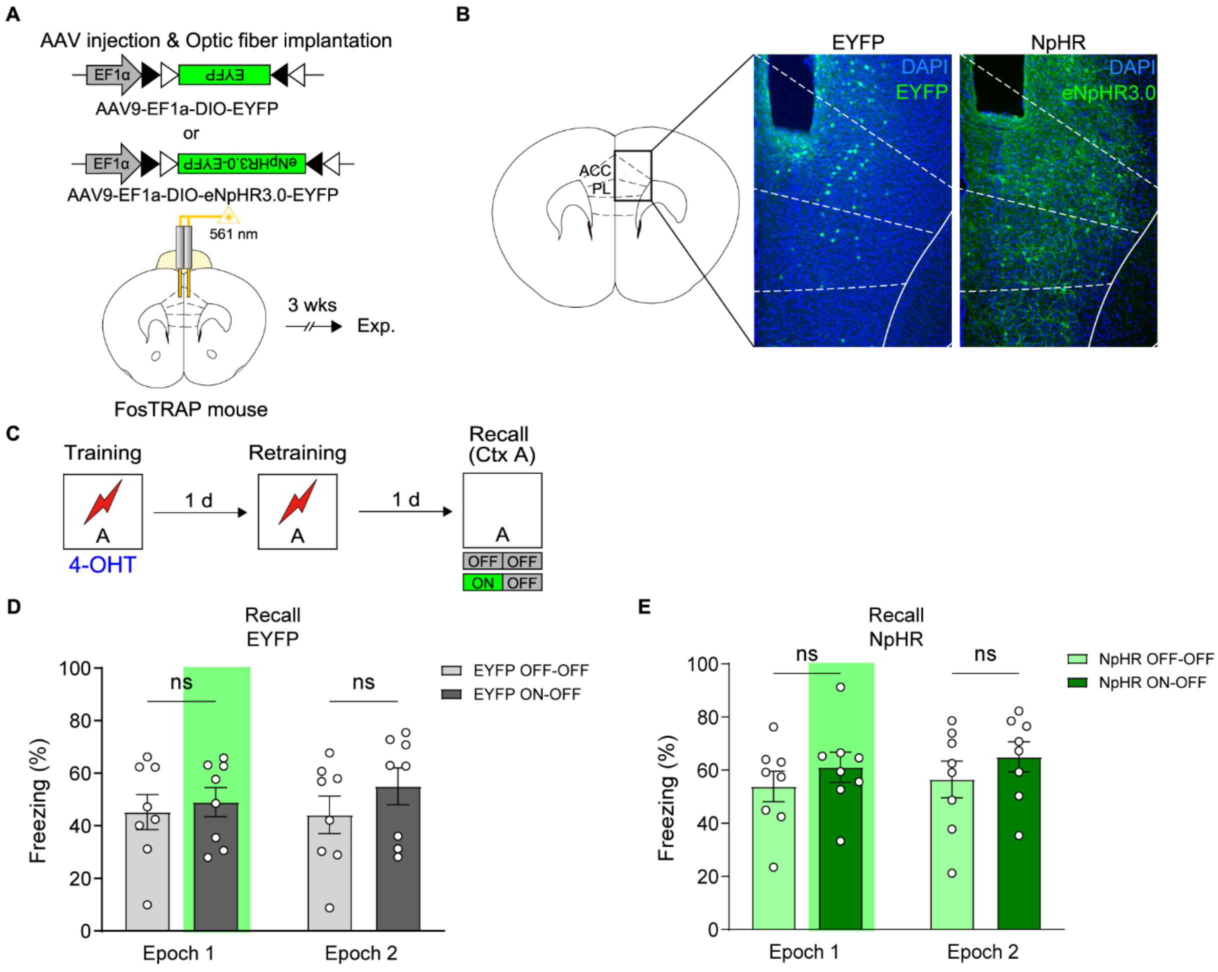


**Figure S6. Initial mPFC c-Fos ensembles are not necessary for memory retrieval in conditioned context**

(A) Schematic of surgery. AAV9-EF1α-DIO-EYFP or AAV9-EF1α-DIO-eNpHR3.0-EYFP was bilaterally injected into the mPFC, followed by implantation of optic fibers above the injection sites. Behavioral experiments were conducted three weeks later.

(B) Representative confocal images of the mPFC from EYFP group (left), and NpHR group (right). Neurons activated during the initial training were labeled with EYFP or eNpHR3.0-EYFP by 4-OHT injection.

(C) Mice underwent initial training followed by 4-OHT injection to label c-Fos–positive ensembles. After retraining, a 6-min context recall test was conducted. Light delivery conditions were divided into ON–OFF (light applied during the first 3 min and absent during the subsequent 3 min) and OFF–OFF (no light during either epoch). Freezing was measured separately during the first 3-min epoch (Epoch 1) and the subsequent 3-min epoch (Epoch 2).

(D) Quantification of freezing behavior during context recall test in EYFP-expressing mice (EYFP; OFF-OFF n=8, ON-OFF n=8). Freezing levels were comparable between the OFF–OFF and ON–OFF groups across both epochs (OFF-OFF vs ON-OFF: p=0.9043 (Epoch 1), p=0.4460 (Epoch 2); Epoch 1 vs Epoch 2: p=0.9587 (OFF-OFF), p=0.2841 (ON-OFF), two-way repeated measures ANOVA with Sidak's multiple comparisons test).

(E) Quantification of freezing behavior during context recall test in NpHR-expressing mice (NpHR; OFF-OFF n=8, ON-OFF n=8). Freezing levels were comparable between the OFF–OFF and ON–OFF groups across both epochs (OFF-OFF vs ON-OFF: $p=0.6456$ (Epoch 1), $p=0.5457$ (Epoch 2); Epoch 1 vs Epoch 2: $p=0.8556$ (OFF-OFF), $p=0.7092$ (ON-OFF), two-way repeated measures ANOVA with Sidak's multiple comparisons test).

Data are shown as mean ± SEM. ns, not significant. See also Table S1.

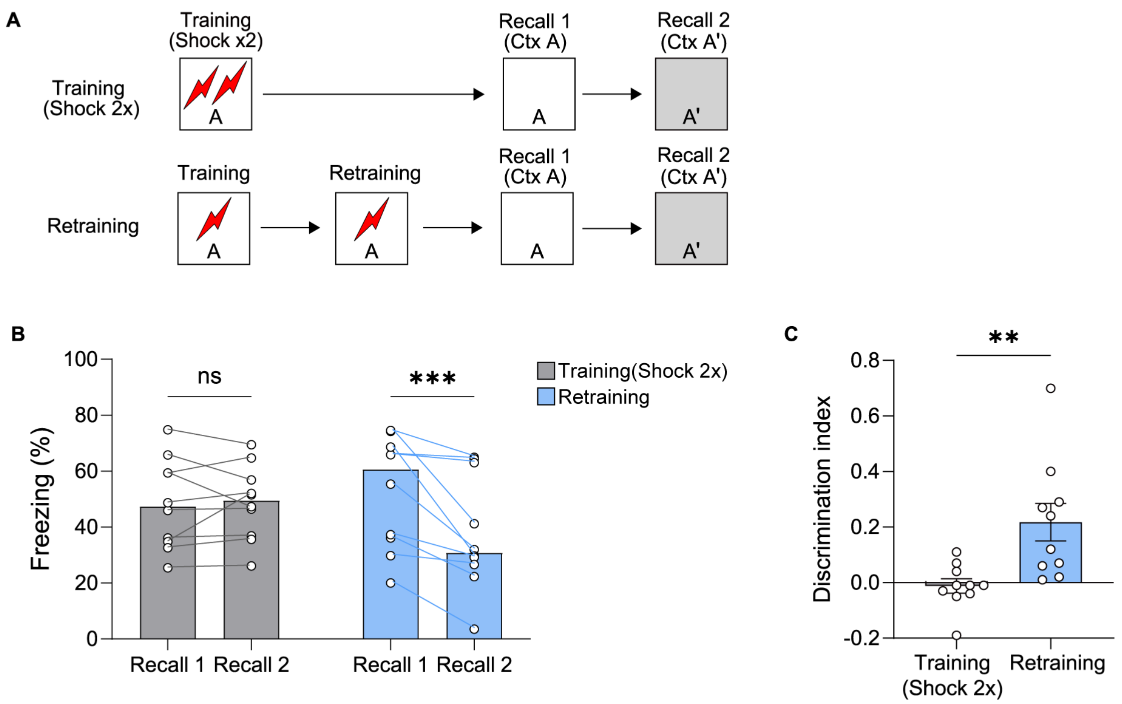


**Figure S7. Context discrimination between the conditioned (Ctx A) and similar (Ctx A') contexts is enhanced by retraining.**

(A) Behavioral procedure. Mice were assigned to either the Training(Shock 2x) group, which received two context–shock pairings, or the Retraining group, which underwent additional training across two consecutive days. Mice were then tested for memory recall in the conditioned context (Recall 1; Ctx A), followed by a recall test in a similar context on subsequent days (Recall 2; Ctx A') (Training(Shock 2x), n = 10; Retraining, n = 10).

(B) Quantification of freezing during Recall 1 (Ctx A) and Recall 2 (Ctx A'). The Training(Shock 2x) group showed comparable freezing levels across the two contexts, whereas the Retraining group displayed a significant reduction in freezing in the similar context (Recall 1 vs Recall 2: p=0.9908 (Training(Shock 2x)), p=0.0007 (Retraining); Training(Shock 2x) vs Retraining: p=0.8207 (Recall 1), p=0.3186 (Recall 2), two-way repeated measures ANOVA with Sidak's multiple comparisons test).

(C) The Retraining group exhibited a significantly higher discrimination index compared to the Training(Shock 2x) group (p=0.0052, unpaired t-test). Discrimination index = (freezing in A − freezing in A')/(freezing in A + freezing in A').

Data are shown as mean ± SEM. ns, not significant. **p<0.01, ***p<0.001. See also Table S1.
